# Combining single-gene-resistant and pyramided cultivars in agricultural landscape compromises the benefits of pyramiding in most, but not all, productions situations

**DOI:** 10.1101/2024.02.14.580232

**Authors:** Marta Zaffaroni, Julien Papaïx, Loup Rimbaud, Abebayehu G. Geffersa, Jean-François Rey, Frédéric Fabre

**Affiliations:** INRAE, Bordeaux Sciences Agro, SAVE, 33882 Villenave d’Ornon, France; INRAE, BioSP, 84914 Avignon, France; INRAE, Pathologie Végétale, 84140 Montfavet, France; CSIRO Agriculture and Food, GPO Box 1700, Canberra, ACT 2601, Australia

**Keywords:** deployment strategy, disease control, durable resistance, evolutionary epidemiology, field size, simulation modelling

## Abstract

**Context:** While resistant cultivars are valuable in safeguarding crops against diseases, they can be rapidly overcome by pathogens. Numerous strategies have been proposed to delay pathogen adaptation (*evolutionary control*), while still ensuring effective protection (*epidemiological control*). Resistance genes can be deployed in 1) single-gene-resistant cultivars sown in the same field (mixture strategy) or in different fields (mosaic strategy), 2) a pyramided cultivar (pyramiding strategy) or 3) hybrid strategies that combine the three previous strategies. In addition, the spatial scale at which resistant cultivars are deployed can affect the plant-pathogens interaction: small fields are thought to reduce pest density and disease transmission.

**Objectives:** We aim to compare these strategies, focusing on the effects of the simultaneous deployment of single-gene-resistant and pyramided cultivars sharing resistance genes in an agricultural landscape. We also investigate the impact of field size.

**Methods:** We used the spatially-explicit stochastic model *landsepi* to compare the evolutionary and epidemiological control across spatial scales and deployment strategies for two major resistance genes.

**Results:** The evolutionary control provided by the pyramiding strategy is at risk when single-gene-resistant cultivars are concurrently planted in the landscape (hybrid strategies). The probabilities of pathogen mutation and the corresponding fitness costs play a crucial role in determining the feasibility of planting pyramided cultivars alongside single-gene-resistant ones. Instead, field size did not affect strategies recommendation.

**Conclusions:** Planting pyramided cultivars alongside single-gene-resistant ones should be avoided. Socio-economic perspectives for the adoption of resistance management strategies are discussed.

## 1 Introduction

Resistant cultivars are widely used in agriculture for the management of pests and diseases. However, their durability is often short lived, with pathogen population rapidly adapting to the deployed resistance genes (McDonald and Linde, 2002; Parlevliet, 2002; Garćıa-Arenal and McDonald, 2003). Several strategies aiming to increase cultivated host genetic diversity have been presented to promote a more durable management of resistant cultivars (McDonald and Linde, 2002; Rimbaud et al., 2021). Increasing the genetic diversity of cultivated hosts challenges pathogens with eco-evolutionary obstacles and delay their adaptation to plant resistance (*evolutionary control*), while providing effective disease protection (*epidemiological control*). For perennial plants, for which rotation is only marginally an option, three basic strategies exist. The first two consist in segregating resistances genes in different single-gene-resistant cultivars and then growing these cultivars together in the same field for “mixture strategy” or in different fields for “mosaic strategy”. Their main expected outcome is to impose disruptive selection on pathogen populations as a result of a heterogeneous host environment favoring different pathogen genotypes at different locations (McDonald and Linde, 2002; Zhan et al., 2015). The third one named “pyramiding strategy”, recently reviewed by Mundt (2018), consists in stacking into a single plant cultivar several resistance genes. Although often costly and time-consuming, breeding pyramided cultivars is a common practice in agriculture often referred by breeders to increase the durability of their varieties. Its main expected outcome is to sharply decrease the probability of the pathogen crossing the mutational pathway that confers adaptation to each resistance gene, known as the “probabilities hypothesis” (Mundt, 1990, 1991).

From an applied perspective, the pyramiding strategy is appealing for several reasons. First, breeders have developed sophisticated approaches to attain resistance pyramids in crops with high agronomic value. Second, implementing pyramided cultivars is easy, as it aligns with the established organization of the agricultural sector between farmers, breeding companies and advisory services, while fitting a widespread expectations in developed agriculture for crop uniformity. By contrast, the deployment of mixture or mosaic of single-gene-resistant cultivars poses some challenges such as the lack of practical rules for their design or the technical concerns for sowing, harvesting and product supply chain (Borg et al., 2018). Third, the history of pyramiding has led to some of the great successes of plant pathology, such as the control of wheat stem rust (Mundt, 2018). From a theoretical perspective, mathematical models suggest that the pyramiding strategy provides a better evolutionary and epidemiological control than mixture and mosaic strategies, for a variety of pathogen reproductive systems, as soon as no adapted pathogens preexist in the pathogen population and mutations rates are low (Lof et al., 2017; Rimbaud et al., 2018a; Zaffaroni et al., 2023). In this setting, the evolutionary control could be achieved for decades while the epidemiological control increases with the proportion of the fields planted with a pyramided cultivar.

Pyramiding strategies present some downsides, as well. First, breeding for pyramiding gets more and more complex and costly as the number of genes to stack increases. By contrast, it is relatively easy to increase the number of single-gene-resistant cultivars deployed in mixture and mosaic strategies. Second, when a pyramided cultivar is overcome by the pathogen, all the involved resistance genes are broken down, hence lost, at once. Theoretical studies suggest that pyramiding can suffer important limitations especially when adapted genotypes are present prior to resistance deployment (Djidjou-Demasse et al., 2017a). Such pre-adapted genotypes, hard to detect when present at low frequencies in the pathogen population, can for example originate from wild relatives of the host cultivar, from which resistance genes are often issued (Garry et al., 2005; Lebeda et al., 2008; Leroy et al., 2014). They can also originate from single-gene-resistant cultivars deployed in the agricultural landscape before the pyramided cultivar (Pink, 2002). This situation may often take place as it is unlikely that breeding programs would forgo the use of resistance genes singly in single-gene-resistant cultivars and wait until these genes can be pyramided (Mundt, 2018). For example, Burdon et al. (2014) reported that the resistant genes against rust that could potentially be stacked in wheat cultivars may have already been used singly in one or more of the breeding programs around the world. Similarly, winegrowers can now plant pyramided cultivars stacking the resistance factors *Rpv1* and *Rpv3* to downy-mildew (Pirrello et al., 2023). If *Rpv1* has never been deployed previously, *Rpv3* was present in many of the interspecific hybrids planted at the beginning of the 20^th^ century. Furthermore, since their approval in 2018 in France, winegrowers can plant single-gene-resistant cultivars bearing *Rpv1* or *Rpv3* along with pyramided cultivars (Merdinoglu et al., 2018). This opens up the possibility of a coexistence in wine-growing areas of single-gene-resistant cultivars bearing *Rpv1* or *Rpv3* with pyramided cultivars bearing both resistance factors.

Despite its practical relevance, the impact of the simultaneous deployment of single-gene-resistant and pyramided cultivars sharing resistance factors in an agricultural landscape has been little studied. In a theoretical study on the evolutionary and demographic dynamics of populations in a source (such as susceptible cultivar) - sink (such as pyramided cultivar) system, the establishment time of a population in the sink habitat resulted to be drastically reduced by the presence of intermediate sinks (such as single-gene-resistant cultivars) (Lavigne et al., 2020). A more realistic, spatially-explicit, modeling study assessed the consequences of the concurrent deployment in an agricultural landscape of two resistance genes singly and pyramided to control epidemics of *P. striiformis* f. sp. *tritici* (Lof et al., 2017). This original work showed that, when virulent pathogen strains have to emerge by mutation (i.e. no preadapted pathogens are initially present), the concurrent use of two single-gene-resistant and one pyramided cultivars favored the emergence of a pathogen breaking down the pyramid. By contrast, the sole deployment of the pyramided cultivar prevented the emergence of a virulent pathogen genotype during the 30 simulated years. These interesting results deserve further investigations. Indeed, the effects of the probability of mutation and of the fitness cost associated to pathogen virulence have not been explored by Lof et al. (2017). Yet, these two parameters, under-investigated in plant pathology, greatly affect the emergence of a super-pathogen adapted to all the deployed resistance genes (Zaffaroni et al., 2023). Moreover, Lof et al. (2017) considered only mosaic strategies where the single-gene-resistant cultivars are deployed in different fields, while these could be planted as mixture in the same field.

Here, we explored the effects of the simultaneous deployment of single-gene-resistant and pyramided cultivars sharing resistance genes in an agricultural landscape. We used the *landsepi* model presented in Rimbaud et al. (2018b) and updated in Zaffaroni et al. (2023), which simulates the spread of epidemics across an agricultural landscape and the evolution of a pathogen in response to the deployment of host resistances. We simulated five strategies involving two major resistance genes. They can be deployed singly in single-gene-resistant cultivars as a mosaic or mixture, or stacked in the same cultivar as a pyramid, that is in “simple strategies”. Single-gene-resistant cultivars and stacked cultivar can also be deployed concurrently within “hybrid strategies” named pyramid-mosaic and pyramid-mixture. In our simulations, we varied two factors related to pathogen epidemiology and evolution (the mutation probability and the cost of virulence) and three factors related to landscape organization (two cropping ratios of fields grown with resistant cultivars and the field size). While we parameterized the model to simulate grapevine downy mildew, which is caused by the oomycete *Plasmopara viticola*, our main conclusions have broader implications to other pathosystems.

## 2 Material and Methods

### 2.1 Model overview

We used the version of the model presented in Zaffaroni et al. (2023). This model simulates the spread and evolution of a pathogen alternating within-season clonal reproduction and between-season sexual reproduction in an heterogeneous agricultural landscape over multiple cropping seasons. The model can handle a range of possible deployment strategies of two major resistance genes (see section 2.2). The demogenetic dynamics of the host-pathogen interaction is based on a compartmental model with a discrete time step, schematically reported in Fig. 1. In the model, H_i,v,t_, L_i,v,p,t_, I_i,v,p,t_, R_i,v,p,t_, and P_i,p,t_ denote the numbers of healthy, latent, infectious and removed individuals, and of pathogen propagules, respectively, in the field *i* = 1,…,J, for cultivar *v* = 1,…, V, pathogen genotype *p* = 1,…,P at time step t=1,…,T*×*Y (Y is the number of cropping seasons and T the number of time steps per season). In this model, an “individual” is defined as a given amount of plant tissue, and is referred to as a “host” hereafter for the sake of simplicity. At the beginning of the first cropping season, susceptible hosts are contaminated with the primary inoculum exclusively composed by non-adapted pathogens (denoted “WT” here for “wild-type”). At the beginning of the following cropping seasons, healthy hosts are contaminated with the primary inoculum generated by a single event of sexual reproduction at the end of the previous cropping season. Propagules sexually produced between cropping seasons are gradually released during the following cropping season. Therefore, within a cropping season, healthy hosts can be infected both by the primary inoculum produced by a single event of sexual reproduction and by the secondary inoculum produced by infected hosts trough multiple events of clonal reproduction. During the simulation, a WT can acquire infectivity gene through a single mutation, or alternatively through sexual reproduction with another pathogen carrying such an infectivity gene. It follows that single mutant “SM1” or “SM2” are able to break down the first or second resistance gene, respectively, and superpathogen “SP” is able to break down both resistance genes (and thus the pyramided cultivar). The acquisition of infectivity, however, may be penalized by a fitness cost *θ* (Leonard, 1977; Brown, 2015), which, in our model, is associated with a lower infection probability. According to Leonard (1977), this fitness cost, could be counterbalanced by the advantage *a* of an adapted pathogen on hosts with corresponding gene of resistance (Table 1). It follows that the adapted pathogen may 1) experience the same fitness cost on any host it can infect, with *θ >* 0 and *a* = 0 (Sapoukhina et al., 2009; Lo Iacono et al., 2012; Nilusmas et al., 2020; Watkinson-Powell et al., 2020; Clin et al., 2021, 2022); 2) experience a fitness cost only for its unnecessary virulence, with *θ* = *a >* 0 (Fabre et al., 2009, 2015; Djidjou-Demasse et al., 2017b; Rimbaud et al., 2018a; Rousseau et al., 2019); 3) not experiencing a fitness cost on any host, with *θ* = 0 and *a* = 0 (Van Den Bosch and Gilligan, 2003; Ohtsuki and Sasaki, 2006; Lo Iacono et al., 2013; Wingen et al., 2013; Lof et al., 2017; Lof and van der Werf, 2017; Pacilly et al., 2018, 2019). For the SP, the fitness costs are multiplicative on every host. The general host-pathogen interaction matrix representing these three cases is reported in Table 1.

**Figure 1:**
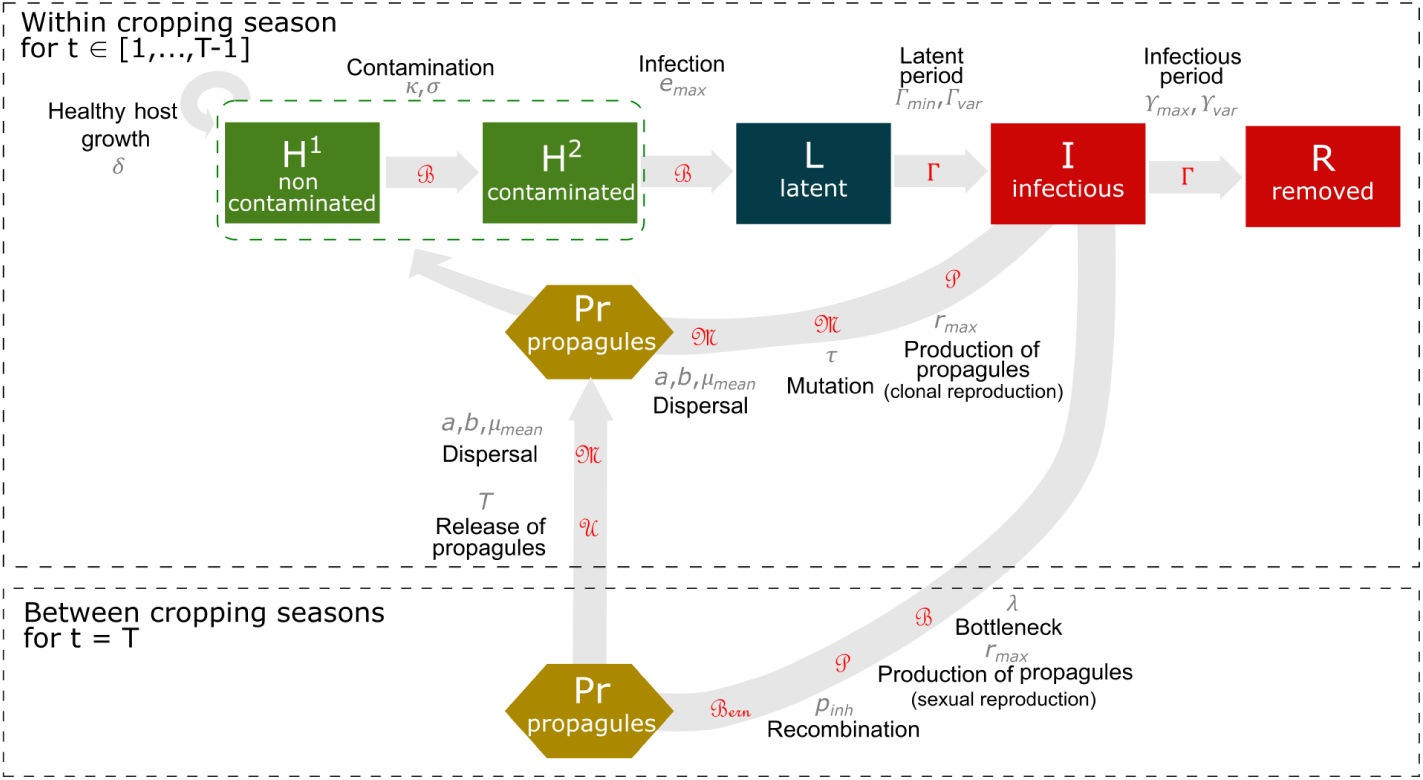
Model overview. Within the cropping season, healthy hosts can be contaminated by pathogen propagules (produced both at the end of the previous cropping season by a single event of sexual reproduction, and within the current cropping season by clonal reproduction) and may become infected. Following a latent period, infectious hosts start producing propagules through clonal reproduction. These propagules may mutate and disperse across the landscape. At the end of the infectious period, infected hosts become epidemiologically inactive. Qualitative resistance prevents the infection of contaminated hosts, *i*.*e*. their transition to the latently infected state. Green boxes indicate healthy hosts contributing to host growth, as opposed to diseased hosts (*i*.*e*. symptomatic, red boxes) or hosts with latent infections (dark blue box). At the end of each cropping season, pathogens experience a bottleneck during the off-season period, and propagules are then produced by sexual reproduction, when genetic recombination may occur. Propagules produced between cropping seasons are gradually released during the following host cropping season. The parameters associated with epidemiological processes are indicated in grey and detailed in Table 2. The distributions used to simulate stochasticity in model transitions are indicated in red; *B*: binomial, Γ: gamma, *P*: Poisson, *M*: multinomial, *U* : uniform, *Bern*: Bernoulli. Host logistic growth is deterministic. Model assumptions and equations are described in *Supporting Information* notes S1 and S2.

**Table 1:** Host-pathogen interaction matrix. The matrix gives the coefficient by which the infection probability is multiplied. The value of this coefficient reflects the relative infection probabilities for the wild-type (WT) and adapted (single mutants SM1 and SM2, and superpathogen SP) pathogen genotypes on the susceptible (SC) and resistant cultivars carrying a single major resistance gene (RC1 and RC2), or their combination (RC12). *θ* is the fitness cost of virulence with respect to the major resistance gene considered and *a* is the advantage of an adapted pathogen on hosts with corresponding gene of resistance.

| | | Host genotype $v$ | | | |
| --- | --- | --- | --- | --- | --- |
|  |  | SC | RC<sub>1</sub> | RC<sub>2</sub> | RC<sub>12</sub> |
| Pathogen genotypes $p$ | WT | 1 | 0 | 0 | 0 |
| | SM<sub>1</sub> | $1-\theta$ | $1-\theta + a$ | 0 | 0 |
| | SM<sub>2</sub> | $1-\theta$ | 0 | $1-\theta + a$ | 0 |
| | SP | $(1-\theta)^2$ | $(1-\theta)(1-\theta + a)$ | $(1-\theta)(1-\theta + a)$ | $(1-\theta + a)^2$ |

**Table 2:** Summary of model parameters and numerical simulation plan (factors in bold are varied according to a complete factorial design).

| Notation | Parameter | Value | Source |
| --- | --- | --- | --- |
| Simulation Factors |  |  |  |
| Y | Number of cropping seasons | 30 years | Fixed |
| T | Number of time steps in a cropping season | 120 days | Fixed |
| V | Number of host cultivars | $[2^a, 3^b, 4^c]$ | Fixed |
| Initial conditions and seasonality (same value for all cultivars) |  |  |  |
| $C_v^0$ | Plantation host density of cultivar v (in pure crops) | 1 m<sup>-2</sup> | Fixed |
| $C_v^{max}$ | Maximal host density of cultivar v (in pure crops) | 20 m<sup>-2</sup> | Fixed |
| $\delta_v$ | Host growth rate of cultivar v | 0.1 day<sup>-1</sup> | [1] |
| $\Phi$ | Initial probability of infection of susceptible hosts | $5 \cdot 10^{-4}$ | Fixed |
| $\lambda$ | Off-season survival probability of pathogen spores | $10^{-4}$ | Fixed |
| Pathogen aggressiveness components |  |  |  |
| $e_{max}$ | Expected infection probability in a compatible host-pathogen interaction | 0.9 | [2,3] |
| $\Gamma_{exp}$ | Expected latent period duration | 7 days | [4] |
| $\Gamma_{var}$ | Variance of the latent period duration | 8 days | [4] |
| $\Upsilon_{exp}$ | Expected infectious period duration | 14 days | [4] |
| $\Upsilon_{var}$ | Variance of infectious period duration | 22 days | [4] |
| $r_{exp}$ | Expected propagule production rate | 2 day<sup>-1</sup> | [4] |
| Sexual reproduction |  |  |  |
| $p_{inh}$ | Probability of a sexual propagule inheriting the genotype at locus $g$ from parent $Par_1$ genotype | 0.5 | Fixed |
| Pathogen dispersal |  |  |  |
| $g(\cdot)$ | Dispersal kernel | Power-law function | See note S2 |
| $\mu_{mean}$ | Mean dispersal distance | 20 m | [4] |
| $\alpha$ | Scale parameter | 5 | Fixed |
| $\beta$ | Width of the tail | 3.5 | [5,6,7] |
| Contamination of healthy hosts |  |  |  |
| $\pi(\cdot)$ | Contamination function | Sigmoid | See note S2 |
| $\sigma$ | Related to the position of the inflection point | 3 | [4] |
| $\kappa$ | Related to the position of the inflection point | 5.33 | [4] |
| Host-pathogen genetic interaction |  |  |  |
| G | Total number of major genes | 2 | Fixed |
| $\tau$ | **Mutation probability** | $[10^{-7}; 10^{-4}]$ | **Varied** |
| $\theta$ | **Fitness cost of virulence** | $[0; 0.05; 0.5]$ | **Varied** |
| $a$ | **Advantage of an adapted pathogen on hosts with corresponding gene of resistance** | $[0; \theta]^d$ | **Varied** |
| Landscape organisation |  |  |  |
| J | **Field size** | **[225; 5625] m<sup>2</sup>** | **Varied** |
|  | **Resistance deployment strategy** | **MI; MO; PY**<br>**PY+MI; PY+MO** | **Varied** |
| $\varphi_1$ | **Cropping ratio of fields where resistance is deployed** | **[0.17; 0.33; 0.5; 0.67; 0.83]** | **Varied** |
| $\varphi_2$ | **Relative cropping ratio of RC<sub>12</sub> in the hybrid strategies** | **[0.33; 0.83]<sup>e</sup>** | **Varied** |

<sup>a</sup> : PY; <sup>b</sup>: MI, MO; <sup>c</sup>: PY + MI, PY + MO; <sup>d</sup>: only for $\theta > 0$ . <sup>e</sup>: only for hybrid strategies. Sources: [1] Bove and Rossi (2020); [2] Bove et al. (2019); [3] Boso and Kassemeyer (2008); [4] Zaffaroni et al. (2023); [5] Frantzen and Van den Bosch (2000); [6] Papaïx et al. (2014); [7] Grosdidier et al. (2018)

Model equations are presented in *Supporting Information*, and the code is available from the R package *landsepi* (v1.3.0, Rimbaud et al. 2023).

#### 2.1.1 Model parameterisation for *Plasmopara viticola*

We parameterised the model to simulate epidemics of *Plasmopara viticola*, the causal agent of grapevine downy mildew, which has a mixed reproduction system: it reproduces clonally during host plant cropping season and sexually at the end of the cropping season (Wong et al., 2001; Gessler et al., 2011). Downy mildew is a major threat to grapevines in all vine-growing areas of the world, causing significant yield losses and leading to the massive use of fungicides (Fouillet et al., 2022). In this context, the commercialization of the new resistant cultivars bearing the resistance genes *Rpv1* and/or *Rpv3* could be a major leverage to reduce treatment and promote agroecological transition as soon as their resistances are not overcome. All the model parameters used in the simulations are listed in Table 2.

### 2.2 Landscape and deployment strategies of resistant cultivars

We considered theoretical agricultural landscapes of 56.25 ha (750 *×* 750 m) composed by regular square-shaped fields (Fig. 2). Two field sizes were considered: small fields (15 *×* 15 m, for an area of 0.025 ha) and large fields (75 *×* 75 m, for an area of 0.5625 ha). Accordingly, the landscape can be composed of 2500 small fields or of 100 large fields. In other words, a single large field is composed by aggregating 25 small fields. In France, the typical vineyard field size range from 0.2 ha in Burgundy vineyard to 0.77 ha in Bordeaux (Adrakey et al., 2022). Up to four cultivars are randomly allocated to fields: a susceptible cultivar (SC) initially infected with a pathogen not adapted to any resistance, two single-gene-resistant cultivars, each carrying a single major resistance gene (RC_1_ and RC_2_), and one pyramided cultivar carrying both resistance genes (RC_12_). The cropping ratio *φ*_1_ represents the proportion of candidate fields in the landscape cultivated with resistant cultivars. For the hybrid strategies combining RC_1_, RC_2_ and RC_12_ we also defined the hybrid cropping ratio *φ*_2_, which represents the relative proportion of fields within the candidate fields cultivated with RC_12_. Candidate fields are planted according to one of the following five strategies, as summarized in Table 3:

1. Mosaics: RC_1_ and RC_2_ are cultivated in equal proportions of the candidate fields;
2. Mixture: both RC_1_ and RC_2_ are cultivated in all the candidate fields, in equal proportions within each field;
3. Pyramiding: RC_12_ is cultivated in all candidate fields.
4. Pyramiding and Mosaic: RC_12_ is cultivated in a proportion *φ*_2_ of the candidate fields, RC_1_ and RC_2_ are cultivated in equal proportions of the remaining 1 *− φ*_2_ proportion of candidate fields;
5. Pyramiding and Mixture: RC_12_ is cultivated in a proportion *φ*_2_ of the candidate fields, RC_1_ and RC_2_ are cultivated in all the remaining 1 *− φ*_2_ proportion of candidate fields, in equal proportions within each field;

**Figure 2:**
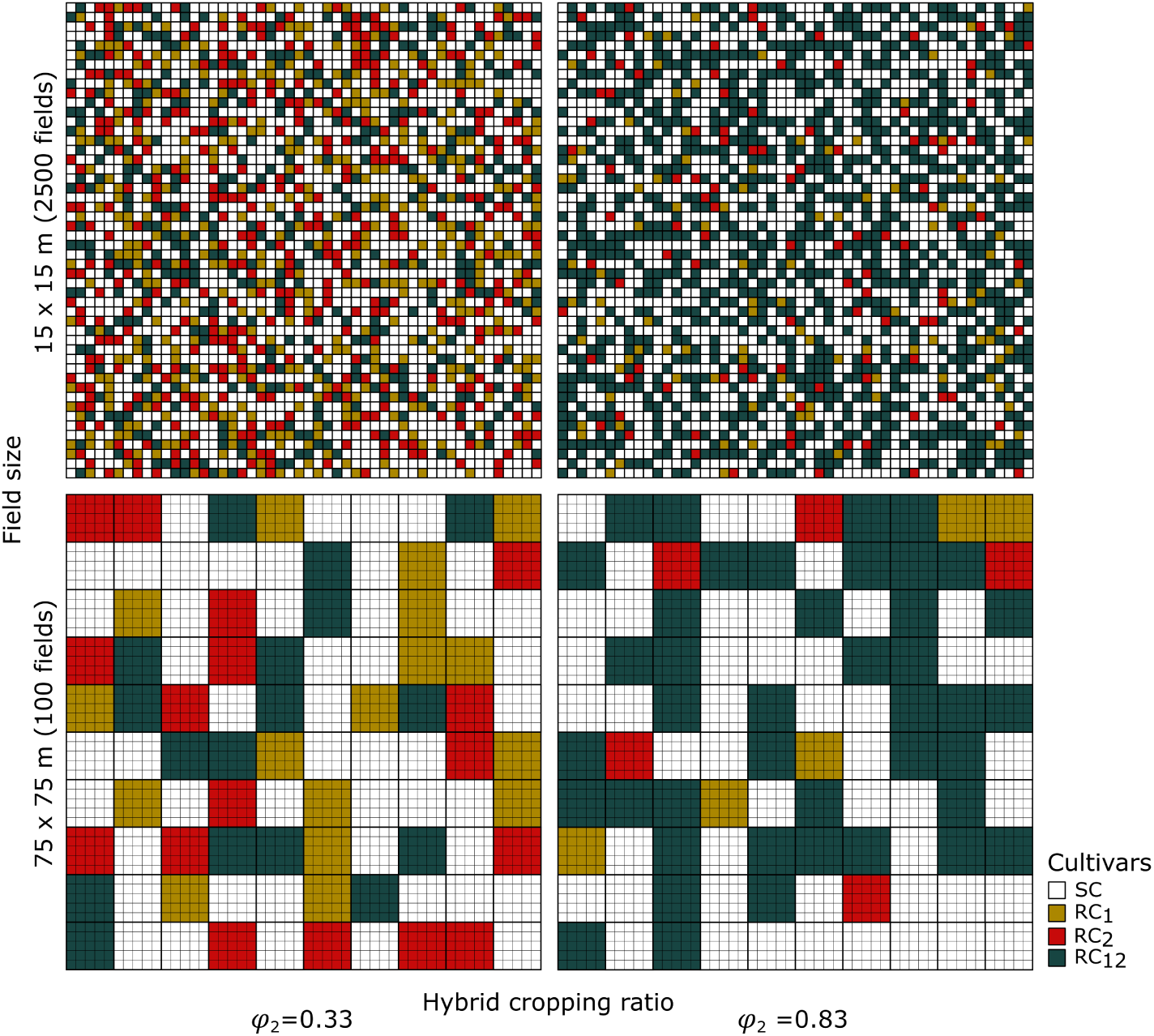
Allocation of four cultivars in the landscape for the hybrid strategy pyramiding and mosaic. In this example, the cropping ratio is fixed at *φ*_1_ = 0.5. The field size varies from small (2500 fields of 15 *×* 15 m) to large (100 fields of 75 *×* 75 m). The hybrid cropping ratio *φ*_2_ varies from 0.33 to 0.83.

**Table 3:**
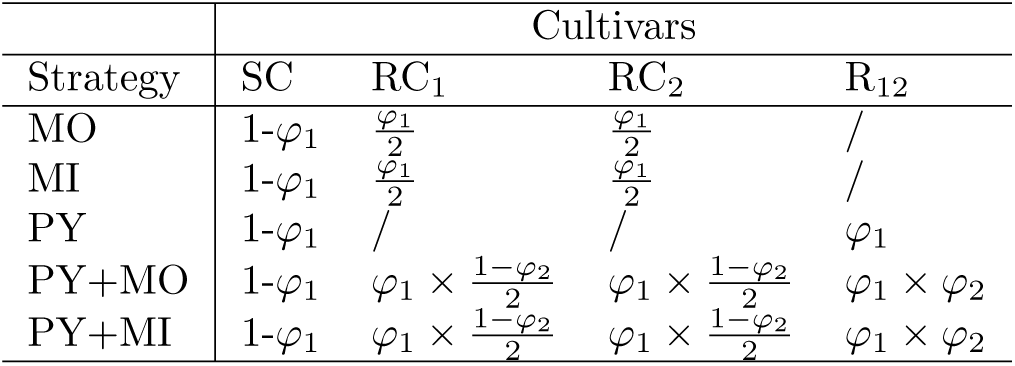
Proportions of fields cultivated with the four cultivars in the considered resistant deployment strategies. *φ*_1_ is the cropping ratio and *φ*_2_ is the hybrid cropping ratio.

| Strategy | Cultivars |  |  |  |
| --- | --- | --- | --- | --- |
|  | SC | RC<sub>1</sub> | RC<sub>2</sub> | R<sub>12</sub> |
| MO | $1-\varphi_1$ | $\frac{\varphi_1}{2}$ | $\frac{\varphi_1}{2}$ | / |
| MI | $1-\varphi_1$ | $\frac{\varphi_1}{2}$ | $\frac{\varphi_1}{2}$ | / |
| PY | $1-\varphi_1$ | / | / | $\varphi_1$ |
| PY+MO | $1-\varphi_1$ | $\varphi_1 \times \frac{1-\varphi_2}{2}$ | $\varphi_1 \times \frac{1-\varphi_2}{2}$ | $\varphi_1 \times \varphi_2$ |
| PY+MI | $1-\varphi_1$ | $\varphi_1 \times \frac{1-\varphi_2}{2}$ | $\varphi_1 \times \frac{1-\varphi_2}{2}$ | $\varphi_1 \times \varphi_2$ |

The remaining proportion 1 *− φ*_1_ of fields are planted with the susceptible cultivar.

### 2.3 Pathogen dispersal

Pathogens disperse within the landscape according to a power-law dispersal kernel, which gives the probability for a pathogen to disperse from one spatial unit to all others in the landscape. By contrast, pathogen dispersal within a spatial unit is assumed homogeneous in space. Here, we consider that a spatial unit is a 15 *×* 15 m square. Hence, one field corresponds to 1 spatial unit in simulations with small fields, and to 25 spatial units in simulations with large fields. It follows that, in the latter, pathogen dispersal is simulated both within and between the 75 *×* 75 m fields.

### 2.4 Simulation plan and model outputs

#### 2.4.1 Simulation plan

The model was used to assess the evolutionary and epidemiological outputs for five deployment strategies (mosaic, mixture, pyramiding, pyramiding and mixture, pyramiding and mosaic). For the simple strategies, we varied the cropping ratio of fields where resistance is deployed (*φ*_1_, five values). We considered different pathogen evolutionary potentials, by varying the mutation probability (*τ*, two levels), the fitness cost of virulence (*θ*, three values) and the way they are paid (*a*, two values). Specifically, for *θ >* 0, we distinguished whether the fitness cost is paid *i*) on any host (*a* = 0) or *ii*) only for unnecessary virulences (*a* = *θ*). For *θ* = 0, we only considered *a* = 0 and this case will be referred to as *θ* = 0 hereafter. Finally, we considered two landscape configurations (small and large fields) as presented in 2.2. The above mentioned factors were explored with a complete factorial design of 300 parameter combinations (Table 2). For the hybrid strategies, in addition to the factors presented above, we varied also the hybrid cropping ratio, *i*.*e*. the relative proportion of candidate fields where the RC_12_ cultivar is planted (*φ*_2_, two values). This results in 400 parameter combinations. Thirty replications were performed for each parameter combination, resulting in a total of 21,000 simulations ((300 + 400) *×* 30). Simulations were run for 30 cropping seasons of 120 days each. Preliminary trials (Zaffaroni et al., 2023) showed that this simulation horizon was sufficiently long to differentiate the performances of the deployment strategies considered. Note that for each simulation the allocation of cultivars to the fields in the landscape was randomly assigned.

#### 2.4.2 Model outputs

At the end of a simulation run, evolutionary and epidemiological outputs were evaluated. For the evolutionary output, we studied SP establishment by defining *E_SP_*, a binary variable set to 1 if the SP becomes established before the end of a simulation run and 0 otherwise. SP is considered established if the number of resistant hosts infected by SP exceeds a threshold above which extinction within a constant and infinite hosts population becomes unlikely. For the epidemiological output, we studied the area under the disease progress curve (AUDPC) to measure disease severity over the whole landscape, averaged across all the simulated cropping seasons. AUDPC is normalised by dividing by the mean disease severity in a fully susceptible landscape; its value therefore ranges from 0 (*i*.*e*. no disease) to 1 (*i*.*e*. disease severity identical to that in a fully susceptible landscape).

### 2.5 Statistical analyses

For each combination of resistance deployment strategy, field size, hybrid cropping ratio, mutation probability, fitness cost and advantage of an adapted pathogen on hosts with corresponding gene of resistance, we fitted second-order logistic or polynomial regressions to assess the response of, respectively, *E_SP_* and AUDPC to variations of cropping ratio. Statistical analyses were performed with the R (v4.0.5, R Core Team 2021) software. The function *geom smooth* within the package *ggplot2* (v3.3.6, Wickham et al. 2022) was used to fit second order logistic regression (method = “glm”, formula = *y ∼* poly(*x,* 2), family = “binomial”) and second-order polynomial regressions (method = “lm”, formula = *y ∼* poly(*x,* 2)).

## 3 Results

Over the 21,000 simulations, the mean AUDPC was 78%, ranging from 13% (2.5th percentile), representing mild epidemics, to 99% (97.5th percentile), representing epidemics almost as severe as in a fully susceptible landscape. The evolutionary output indicated that the superpathogen SP became established before the end of the 30-year simulation in 71% (14,924) of the simulations. Below, we detail how such variability in outputs is driven by the deployment strategies and pathogen features, focusing on the landscape with large fields. Indeed, field size did not (or only marginally) impact the conclusions of the analyses (Fig. S1-S2-S3).

### 3.1 Comparative performance of simple versus hybrid deployment strategies

The mutation probability and the scenarios on fitness costs are a major driver of the performances of the strategies. At low mutation probability and for null fitness cost (*θ* = 0), simple strategies better prevented SP establishment than hybrid strategies (Fig. 3A). In particular, the SP never got established for PY, while for MI and MO, it got established at least in one over two simulations. By contrast, the SP got established in all the simulations involving hybrid strategies. Similarly, when the adapted pathogen experienced a high fitness cost only for its unnecessary virulence (*θ* = 0.5 and *a* = 0.5), simple strategies better prevented SP establishment than hybrid strategies. As previously, the SP never got established for PY. The probability of SP establishment ranged between 0.12 and 0.80 for MI and MO and between 0.30 and 1 for hybrid strategies. In sharp contrast, when the adapted pathogen experienced a high fitness cost on any host (*θ* = 0.5 and *a* = 0), there were no differences between simple and hybrid strategies, all of them limiting the establishment of the SP, with probabilities of SP establishment always lower than 0.15.

**Figure 3:**
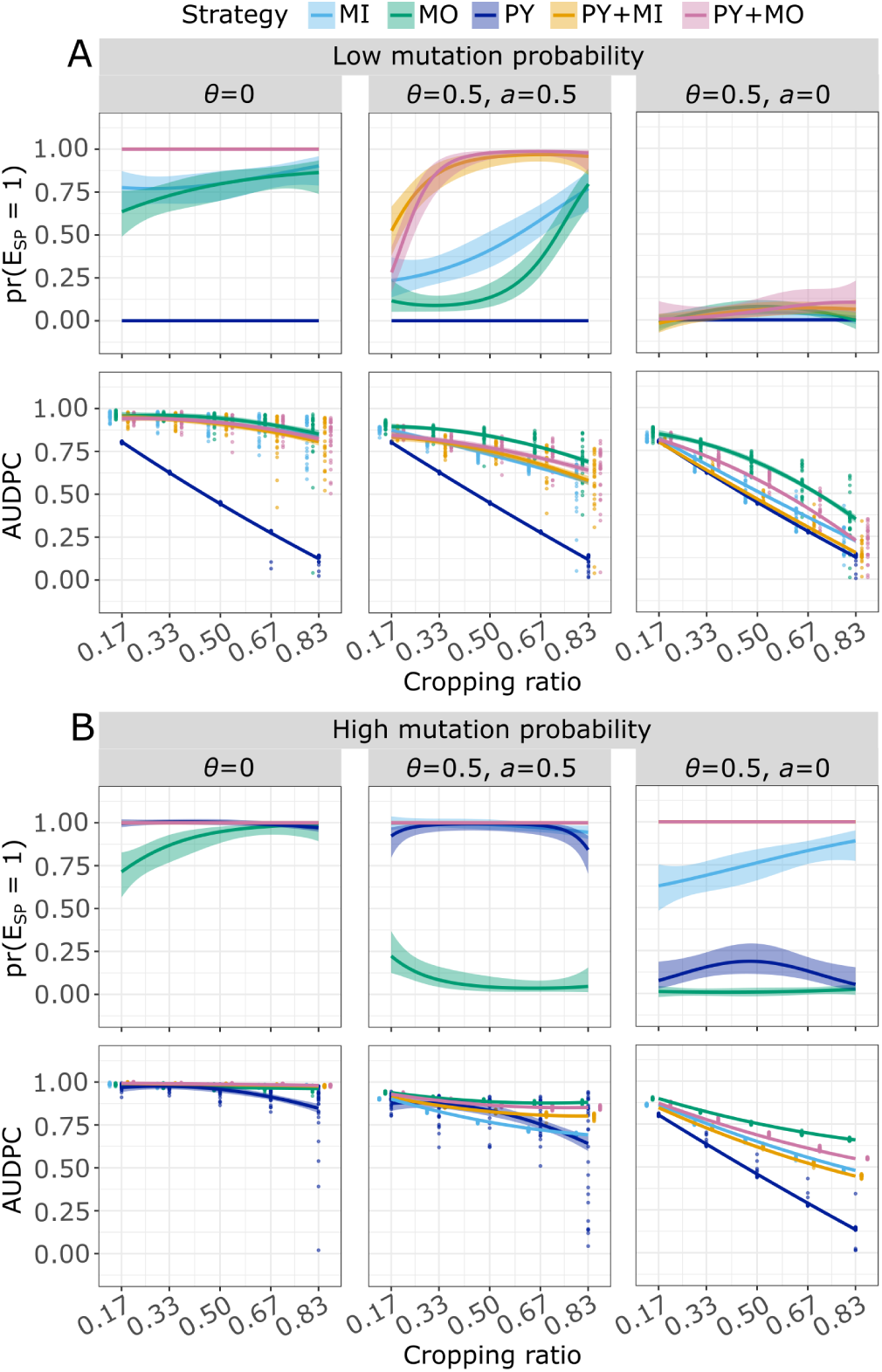
Probability of SP establishment (first row of each panel) and AUDPC (second row of each panel) at low (A, *τ* = 10*^−^*^7^) and high (B, *τ* = 10*^−^*^4^) mutation probability, with no fitness cost (*θ* = 0), with fitness cost for unnecessary virulence (*θ* = 0.5 and *a* = 0.5), or with fitness cost experienced on any host (*θ* = 0.5 and *a* = 0). Simulations were run with the landscape with 100 large fields. Panels show the effect on the probability of *E_SP_* and AUDPC as a function of the cropping ratio for the five resistant deployment strategies considered. Curves are based on the fitting of second-order logistic (first row) or polynomial regressions (second row) to simulation outputs (represented by points, note that, in the first row, we omitted them, being either 0 or 1); shaded envelopes delimited the 5th and 95th percentiles. When the curves or the shaded envelopes extend beyond the 0-1 range for the probability of *E_SP_*, it indicates that fitting a second-order logistic regression was impossible, thus a second-order polynomial regression was fitted instead. When the curve for PY+MO is not visible, it overlaps with that of PY+MI.

With respect to the epidemiological control, simple strategies were not consistently better than hybrid strategies. On the one hand, PY always provided the best epidemiological control, reducing AUDPC by 87% at the highest cropping ratio, independently of the scenario for the type of fitness costs. This resulted from the absence of SP establishment. On the other hand, MO always provided the worst epidemiological control, reducing the AUDPC up to 16% (*θ* = 0), 33% (*θ* = 0.5 and *a* = 0.5) and 66% (*θ* = 0.5 and *a* = 0). The epidemiological control provided by MI, PY+MI and PY+MO laid between these two contrasted situations depending on the fitness costs. For null fitness cost, the performance of MI, PY+MI and PY+MO were close to each other as well as to MO. The situation was the same for high fitness cost paid only for unnecessary virulence even though these three strategies had a slight advantage over MO (reducing the AUDPC by 7% respect to MO). In these two settings, PY always provided a much better epidemiological control. This was no longer true for high fitness cost paid on any host. The epidemiological control of MI, PY+MO and especially of PY+MI were then relatively close to the one provided by PY. The performance of MO was further behind in this setting.

These results were substantially modified at high mutation probability. However, hybrid strategies remained the worst strategies regarding the probability of SP establishment. The SP certainly got established at any cropping ratio for the three scenarios on fitness costs with hybrid strategies. By contrast, the propensity of the simple strategies to prevent SP establishment changed drastically under these three scenarios. In details, without fitness cost, the SP certainly got established at any cropping ratio (Fig. 3B) for PY and MI while the probability of SP establishment rapidly increased to 1 with MO as soon as the cropping ratio exceed 0.5. With high fitness cost paid only for unnecessary virulence, the SP almost certainly got established at any cropping ratio (Fig. 3B) for PY and MI while the probability of SP establishment rapidly decreases to 0 with MO as soon as the cropping ratio exceed 0.5. Finally, for high fitness cost paid on any host, the SP establishment was uncertain for MI, with associated probability increasing from 0.67 to 0.9 with the cropping ratio. This probability never exceed 0.25 for PY and was almost 0 for MO.

The epidemiological control was worsened at high mutation probability, in particular for PY even if this strategy generally remained the best. Without fitness cost and for cropping ratio lower than 0.5, the epidemiological control was weak (with AUDPC at best reduced by 3%) whatever the strategies considered. At higher cropping ratio, PY was the only strategy that stood out with AUDPC reduction up to 16% at the highest cropping ratio of 0.83. With high fitness cost paid only for unnecessary virulence, simple strategies PY and MI better controlled the epidemics than hybrid strategies and, notably, MI slightly better performed than PY for intermediate cropping ratio. The highest average AUDPC reduction achieved for PY was 37% at cropping ratio of 0.83, a value nearly identical to the epidemiological control provided by MI. Hybrid strategies were rolled back, with AUDPC reduction up to 15% and 20%, respectively for PY+MO and PY+MI. As for low mutation probabilities, MO provided the worst epidemiological control, reducing the AUDPC up to only 12%. For high fitness cost paid on any host, PY regained a clear advantage over the four others strategies, with a reduction of the AUDPC up to 86%, a value nearly identical to the level of control achieved at low mutation. The simple strategy MI provided almost the same epidemiological control than its hybrid counterpart PY+MI, with AUDPC reductions up to 52% (for MI) and 55% (for PY+MI). By contrast, the hybrid strategy PY+MO provided a better epidemiological control than the simple strategy MO, reducing the AUDPC respectively up to 45% and 34%.

### 3.2 Effect of the hybrid cropping ratio on the performance of hybrid versus pyramid strategies

In the hybrid strategies analyzed previously, the relative proportion of pyramided cultivar *φ*_2_ was set to 0.33 meaning that the cultivars *RC*_1_, *RC*_2_ and *RC*_12_ all occupied the same surface in the landscape. A further question was to which extend the performance of hybrid strategies would improve at higher *φ*_2_. We compared below relative proportions of pyramided cultivar of 0.33 and 0.83. At low mutation probability, increasing *φ*_2_ from 0.33 to 0.83 decreased the probability of SP establishment only when adapted pathogen experienced high fitness cost for its unnecessary virulence (Fig. 4, *θ* = 0.5 and *a* = 0.5). Few changes were obtained with the other two scenarios on fitness costs (*θ* = 0 or *θ* = 0.5 and *a* = 0). However, increasing *φ*_2_ substantially improved the epidemiological control both without fitness cost (*θ* = 0) and with high fitness costs paid only for unnecessary virulence (*θ* = 0.5 and *a* = 0.5) . Specifically for the PY+MI strategy, at the highest cropping ratio and fitness cost, the average AUDPC reduction increased from 43% (*φ*_2_ = 0.33) to 68% (*φ*_2_ = 0.83). Similarly, for the PY+MO strategy, the average AUDPC reduction increased from 37% (*φ*_2_ = 0.33) to 60% (*φ*_2_ = 0.83). However, the level of control obtained remained far behind the 88% AUDPC reduction obtained when deploying the pyramided cultivar only. With high fitness cost paid on any host (*θ* = 0.5 and *a* = 0), increasing *φ*_2_ exclusively improved the epidemiological control provided by the PY+MO strategy. Specifically, at the highest cropping ratio, the average AUDPC reduction increased from 78% (*φ*_2_ = 0.33) to 86% (*φ*_2_ = 0.83).

**Figure 4:**
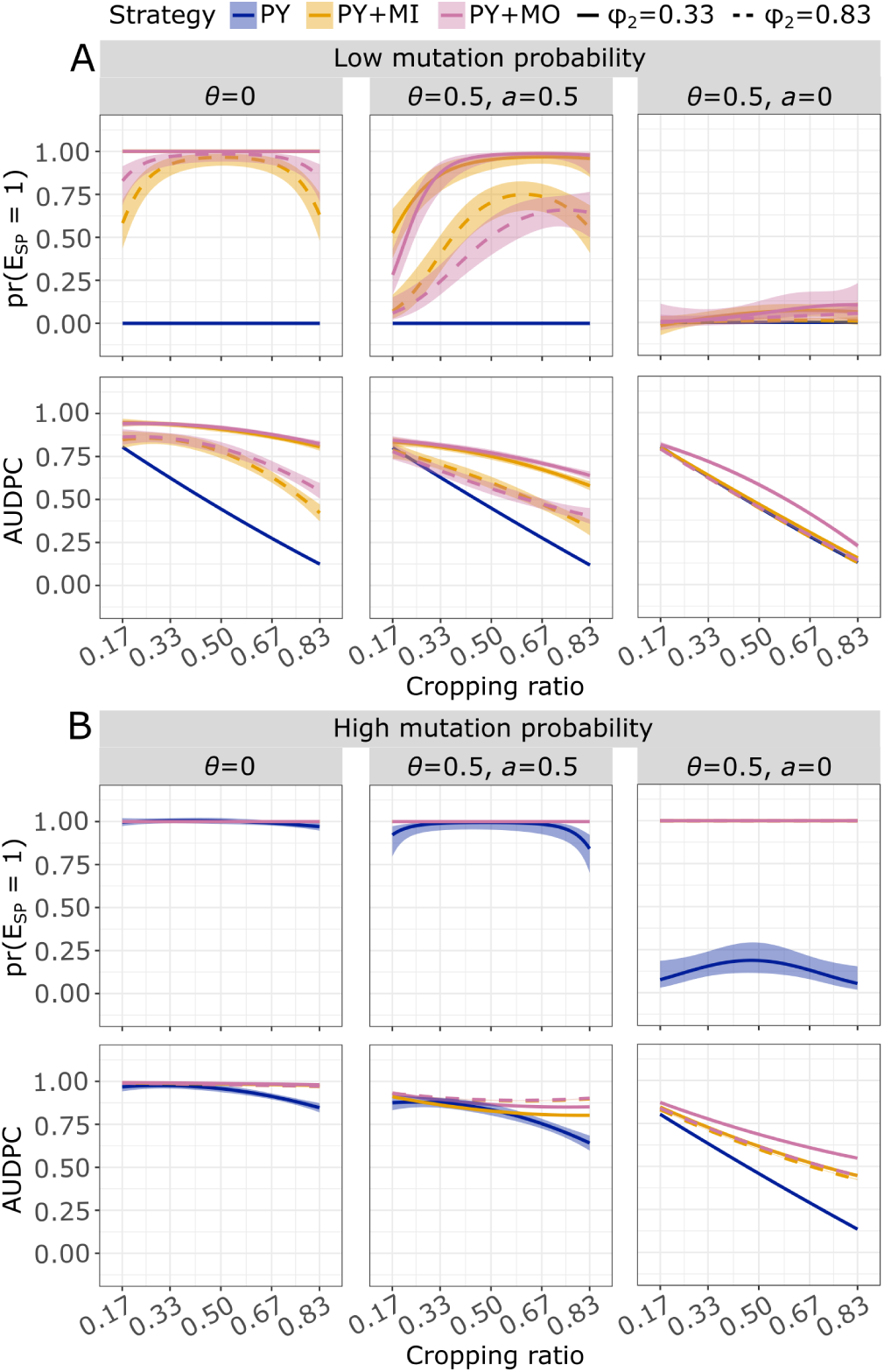
Probability of SP establishment (first row of each panel) and AUDPC (second row of each panel) at low (A, *τ* = 10*^−^*^7^) and high (B, *τ* = 10*^−^*^4^) mutation probability, with no fitness cost (*θ* = 0), with fitness cost for unnecessary virulence (*θ* = 0.5 and *a* = 0.5), or with fitness cost experienced on any host (*θ* = 0.5 and *a* = 0). Simulations were run with the landscape with 100 large fields. Panels show the effect on the probability of *E_SP_* and AUDPC as a function of the cropping ratio for pyramiding and for the hybrid strategies, at different proportions of the pyramided cultivars with respect to single-generesistant cultivars (*φ*_2_). Curves are based on the fitting of second-order logistic (first row) or polynomial regressions (second row) to simulation outputs; shaded envelopes delimited by the 5th and 95th percentiles. When the curves or the shaded envelopes extend beyond the 0-1 range for the probability of *E_SP_*, it indicates that fitting a second-order logistic regression was impossible, thus a second-order polynomial regression was fitted instead.

Very limited effects of increasing *φ*_2_ were observed at high mutation probability for both the evolutionary and the epidemiological control. The only noticeable effect was an improvement of the epidemiological control for PY+MO strategy, under the scenario that high fitness cost was paid on any host. Specifically, at the highest cropping ratio, the average AUDPC reduction increased from 45% (*φ*_2_ = 0.33) to 55% (*φ*_2_ = 0.83).

### 3.3 Effect of intermediate fitness cost on the performance of hybrid strategies

Previous results suggested that the performance of the strategies considered, and particularly the performance of hybrid strategies compared to PY, were strongly impacted by whether high fitness cost (*θ* = 0.5) were paid only for unnecessary virulence or on any host. Specifically, PY over-performed the hybrid strategies PY+MI and PY+MO (*i*) at low mutation probability when a high fitness cost was paid only for unnecessary virulence and (*ii*) at high mutation probability when a high fitness cost was paid on any host. (Fig. 5). To go a step further, we now considered a lower fitness cost (*θ* = 0.05). The comparison between hybrid strategies and MI and MO can be found in the *Supporting Information* (Fig. S4-S7).

**Figure 5:**
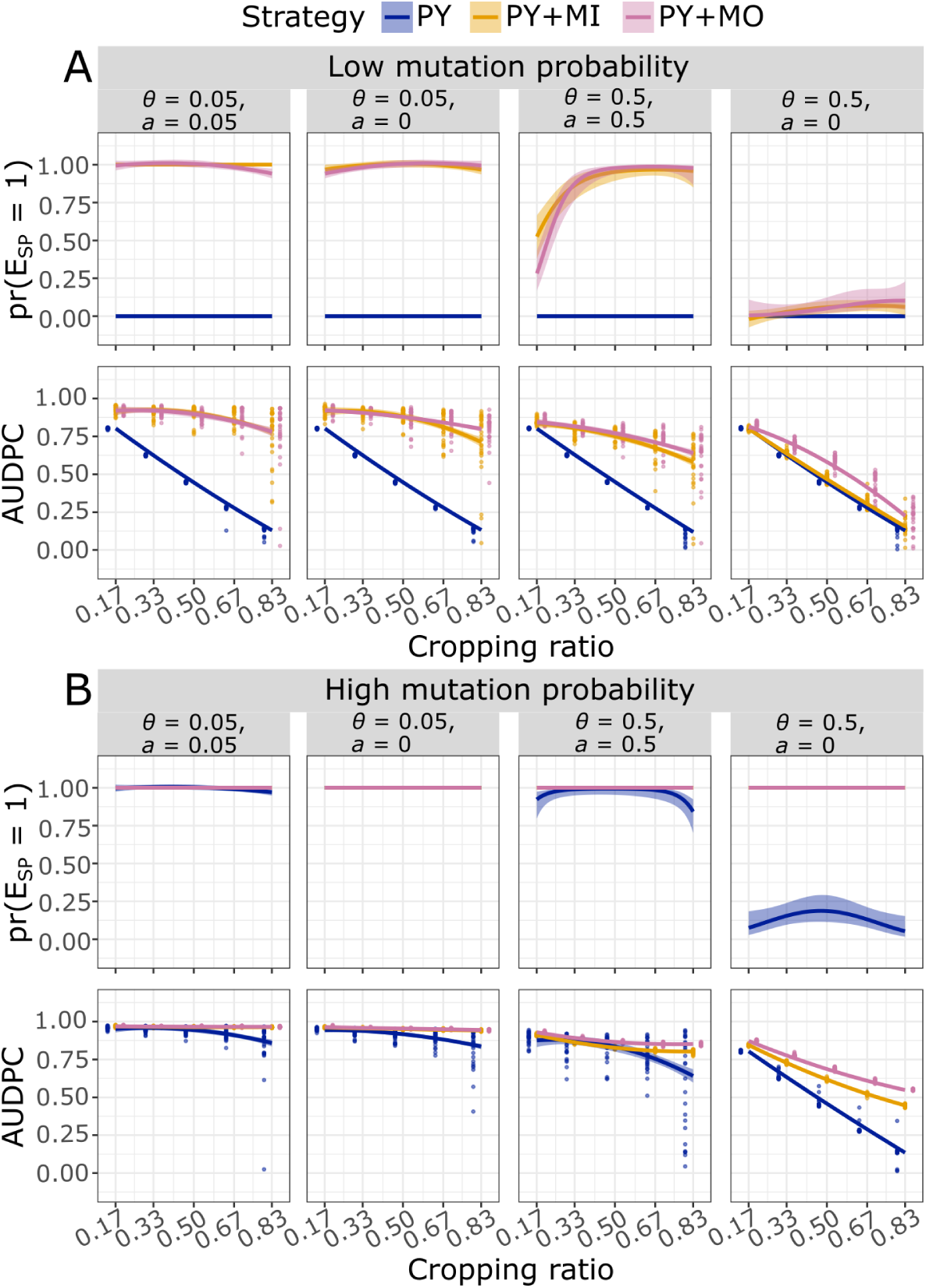
Probability of SP establishment (first row of each panel) and AUDPC (second row of each panel) at low (A, *τ* = 10*^−^*^7^) and high (B, *τ* = 10*^−^*^4^) mutation probability, with intermediate or high fitness cost (resp. *θ* = 0.05 and *θ* = 0.5), which is paid for unnecessary virulence (resp. *a* = 0.05 and *a* = 0.5), or on any host (*a* = 0). Simulations were run with the landscape with 100 large fields. Panels show the effect on the probability of *E_SP_* and AUDPC as a function of the cropping ratio for pyramiding and for the hybrid strategies. Curves are based on the fitting of second-order logistic (first row) or polynomial regressions (second row) to simulation outputs; shaded envelopes delimited by the 5th and 95th percentiles. When the curves or the shaded envelopes extend beyond the 0-1 range for the probability of *E_SP_*, it indicates that fitting a second-order logistic regression was impossible, thus a second-order polynomial regression was fitted instead. When the curve for PY+MO is not visible, it overlaps with that of PY+MI.

In sharp contrast with the scenarios at high fitness cost, no more (or very few) differences exist between hybrid and PY strategies in presence of an intermediate cost between the scenarios where it is paid only for unnecessary virulence (*θ* = 0.05 and *a* = 0.05) or on any host (*θ* = 0.05 and *a* = 0, Fig. 5). Moreover, the evolutionary and epidemiological performance obtained with intermediate cost under both scenarios are quite close to the one obtained with high fitness costs paid only for unnecessary virulence (*θ* = 0.5 and *a* = 0.5). At high mutation probability, hybrid and PY strategies shared similar performances in these three scenarios (Fig. 5B).

At low mutation probability, the evolutionary control was always better for PY than for hybrid strategies. The epidemiological control was also better for PY than for hybrid strategies, the over-performance of PY increasing linearly with the cropping ratio. Finally, our results showed few differences between the two hybrid strategies PY+MI and PY+MO considered when compared to PY.

## 4 Discussion

### 4.1 Pyramided cultivars may be at risk when single-gene-resistant cultivars are planted at the same time

The superiority of pyramiding among the strategies considered is the result of the multiple mutations required for the emergence of a superpathogen adapted to the stacked resistance genes (SP). Either these mutations must be acquired at the same time, but the probability of such an event is weak, or stepwise. The latter case requires the existence of single mutants pathogens that are: 1) possibly less competitive than wild type ones on susceptible hosts and 2) not able to infect the pyramided cultivar. This explains why, at low mutation probability or high fitness cost of virulence, pyramiding has been proven to be the best strategy in controlling the epidemic and avoiding the breakdown of resistance genes (Lof et al., 2017; Rimbaud et al., 2018b; Zaffaroni et al., 2023). However, our results indicated that concurrently deploying a pyramided cultivar along with single-gene-resistant cultivars, carrying the resistance genes stacked into the pyramid, could often result in a drastic decrease in the evolutionary and epidemiological control provided by pyramiding. This is the case when the adapted pathogen does not experience a fitness cost on any host (*θ* = 0), or when it experiences such cost exclusively for its unnecessary virulence (*θ* = 0.5 and *a* = 0.5). Indeed, in these situations, single-gene-resistant cultivars, even planted in small proportion, can easily select single mutants that serve as stepping stones for the emergence of the SP. By contrast, when the adapted pathogen experiences a high fitness cost on any host in the landscape (*θ* = 0.5 and *a* = 0), pathogen evolution is slowed down. It takes longer for the single mutant to establish as it pays a fitness cost on all the hosts in the landscape. Consequently, single-gene-resistant cultivars are less likely to act as stepping stones for the SP emergence, and the establishment of the SP is no longer influenced by planting pyramided cultivars alongside single-resistant ones (the mutation probability being too low to see an effect, see further). In that situation, hybrid strategies, in particular the one combining pyramiding and mixture, provides evolutionary and epidemiological control similar to that provided by pyramiding alone.

For a pathogen with high mutation probability, the superiority of pyramiding over the other deployment strategies is greatly reduced in the absence of fitness cost or with cost only for unnecessary virulence. In this setting, the SP almost certainly established for strategies involving pyramided cultivars. Consequently, concurrently deploying single-gene-resistant and pyramided cultivars only marginally reduces the evolutionary and epidemiological controls provided by simple pyramiding strategy. In sharp contrast, pyramiding maintains its superiority over hybrid strategies when adapted pathogen experiences a high fitness cost on any host. Remarkably, it should also be noted that mixture could be preferred over pyramiding at high cropping ratio and when fitness cost is paid on any host.

Overall, our results are coherent with those of Lof and colleagues (Lof et al., 2017; Lof and van der Werf, 2017). They showed that, without fitness cost, concurrent deploying single-gene-resistant and pyramided cultivars greatly decreased the durability of the pyramided cultivar when the SP must appear by stepwise mutations, that is when pre-adapted pathogen are initially absent from the population. Our work goes a step beyond as we considered that adapted pathogen can experience a fitness cost on any host or exclusively for its unnecessary virulence. In previous modeling studies, authors followed either the first (Sapoukhina et al., 2009; Lo Iacono et al., 2012; Nilusmas et al., 2020; Watkinson-Powell et al., 2020; Clin et al., 2021, 2022) or the second approach (Fabre et al., 2009, 2015; Djidjou-Demasse et al., 2017b; Rimbaud et al., 2018a; Rousseau et al., 2019). Here, for the first time, we compared the two evolutionary scenarios, shading light on the implications of the structure of the host-pathogen interaction matrix on the performance of simple *vs*. hybrid resistant deployment strategies. Our results showed that, when the fitness cost is high, the structure of the host-pathogen interaction matrix is a key factor determining whether or not pyramiding loses its efficacy when concurrently deployed with single-gene-resistant cultivars. By contrast, for intermediate fitness cost, the performance of simple *vs*. hybrid strategies is not affected by the structure of the host-pathogen interaction matrix. This is of practical importance as empirical detection of measurable fitness costs has often been difficult or impossible for cellular plant pathogen, which could suggest that such costs can be low (Leach et al., 2001; Sacristán and Garćıa-Arenal, 2008). In this work, we considered only two cases along the continuum of plant-pathogen interactions spanning from pure gene-for-gene (*θ ≥* 0 and *a* = 0) to matching allele interactions (*θ* = 1 and *a* = 1) (Agrawal and Lively, 2002; Mikaberidze et al., 2015). Continuously varying *θ* and *a* will allow to further explore the effects of this continuum of interactions using the same framework (Table 1). From an applied perspective, our results suggested that practitioners aiming to guide the choice of a deployment strategy for a given pathosystem have to carry out extensive experimental work informing the fine structure of the host-pathogen interaction matrix.

### 4.2 Policies to favor the durability of pyramiding

Without accurate knowledge on the host-pathogen interaction matrix, it is reasonable to avoid planting pyramided cultivars alongside single-gene-resistant ones. Putting into practice this rule is however challenging due to the lack of legal regulations on this issue and the existence of many barriers to change current habits. These barriers result from the interaction between the social, political, and institutional actors involved and are reinforced by the spatial scale (region, production area) at which management strategies must be designed.

Managing resistance genes, akin to finite natural resources, demands careful stewardship, involving dedicated policies and incentives. To design these actions, it is first important to stress that effective resistance management strategies provide substantial economic benefits across scales and stakeholders. For farmers, durable genetic resistance provides direct benefits, such as improved and consistent quantitative and qualitative yields. Additionally, they experience indirect benefits, including decreased reliance on pesticides, simplified pest management, working time savings for implementing other valuable on-farm operations, and the possibility to redirect financial resources previously allocated to pesticides and dedicated spreaders towards enhancing various production activities (Geffersa et al., 2023). For breeding companies, durable genetic resistance can generate spillover benefits by freeing resources otherwise needed to deal with pest evolution and pesticide resistance. These resources could for example be redirected to new R&D investments such as improving agronomic cultivar traits favoring the adoption of sustainable practices. Finally, at a broader scale, durable genetic resistance provides indirect benefits by reducing chemical reliance, cutting health costs and boosting export demand by minimizing chemical residues (Kahindo and Blancard, 2022; Abedullah et al., 2015; Pimentel et al., 1992).

The diversity of stakeholders involved (farmers, breeders, consumers, industry, researchers, practitioners, and policymakers) combined to their own costs and benefits lead to highly contrasted perceived value of investing in stewardship (Geffersa et al., 2023). This diversity of externalities implies that the efforts of one stakeholder to preserve resistance durability is likely to have very little effect if the others do not cooperate (Ambec and Desquilbet, 2012). This is especially the case with highly mobile plant pathogens dispersing with airborne propagules. The efforts done by a few farmers to preserve resistance durability could be useless since adapted pathogen can emerge and spread from nearby fields managed by not so forward-looking farmers. A strong collaborative approaches is then needed for the implementation of successful stewardship strategies, as already shown for pesticides resistance management (Lazarus and Dixon, 1984; Lemaríe and Marcoul, 2018). A growing number of case studies have identified factors that impede the success of institutional arrangements and collaborative efforts in the context of sustainable agriculture (Fischer et al., 2019; Leventon et al., 2017; Velten et al., 2021). This underscores a need for increased government efforts to facilitate collaboration and coordination in stewardship between farmers and multiple stakeholders to support the harmonization of multiple objectives through a comprehensive approach integrating education, incentives, legislation, and social change for sustainable genetic control of crop diseases and resistance durability (Geffersa et al., 2023). How to best integrate regulation, pecuniary economic incentives and non-pecuniary instruments have been deeply modeled and analyzed for proposing optimal policy ensuring both efficiency and durability of GM Bt crop in US and in Australia (Bourguet et al., 2005; Vacher et al., 2006; Brown, 2018; Smulders et al., 2021).

Among the various regional-scale policies proposed to control pathogen spread and increase resistance durability, one option currently discussed is diversifying cropland and reducing field size on a landscape level (Tscharntke et al., 2021). Recently, Sirami et al. (2019) evidenced over 435 agricultural landscapes located in 8 European and North American regions that increasing configurational cropland heterogeneity by decreasing field size can be as beneficial for multitrophic diversity (plants, birds, bees, butterflies, carabid beetles, spiders, and syrphid flies) as increasing seminatural habitat. Here, we tested whether the effect of decreasing field area by 25 impacted the strategy recommendations for various fitness costs, mutation probabilities and cropping ratios. Minor, if any, effects were evidenced. In particular disease severity was only marginally affected by the field size (Fig. S1-2 in the *Supporting Information*). This is due, for example, to the relatively high proportions (0.45 for small and 0.82 for field) of infectious propagules remaining in the same field they were produced. Ultimately, this favours the dilution effect provided by mixture over mosaic. Any factor lowering the value of this proportion (such as larger pathogen dispersal distances), is likely to improve the dilution effect provided by mosaic.

### 4.3 Further perspective

In our work, we considered that the fitness costs remained the same for the 30 years of simulation. In reality, compensatory mutations can arise in the genome at secondary loci following resistance breakdown, possibly compensating the cost of virulence (Bahri et al., 2009). Compensatory mutations can thus impact the outcomes of our study, as virulent pathogens would no longer pay a high fitness cost in the long run. For example, at high mutation probability and for a fitness cost paid on any host, compensatory mutations could nullify the advantages of deploying pyramiding alone compared to hybrid strategies (Fig. 5B). It would be interesting to integrate this possibility in the model and explore its effects on short and long-term epidemiological and evolutionary dynamics (Day and Gandon, 2012). Furthermore, we considered here only qualitative resistance genes (i.e. major genes), although quantitative resistance is gaining interest for use in pathogen control (Niks et al., 2015; Parlevliet, 2002). Quantitative resistance genes are thought to be more durable than qualitative ones, as a result of the accumulated effect of several minor-effect genes (Niks et al., 2015). As the model *landsepi* can also handle quantitative resistances, it would be intriguing to expand our analysis in this aspect, for example considering strategies involving both qualitative and quantitative resistance genes. Rimbaud et al. (2018b) showed that pyramiding a qualitative and a highly efficient quantitative resistance gene, rather than two qualitative genes, increased the durability of the strategy. Fabre et al. (2022) showed that the choice of the quantitative resistance genes operated by breeders, throughout their differential effects on pathogen traits, differential impacted pathogen diversification. How these results could be impacted when considering hybrid strategies based on combination of quantiative and qualitative resistances remained an open question.

## Acknowledgments

We are grateful to Stephane Lemarié (INRAE) for his explanations on economics of innovation in agriculture.

## Statements & Declarations

### Author contributions

### Data availability

## References

Abedullah, Kouser, S., and Qaim, M. (2015). Bt cotton, pesticide use and environmental efficiency in pakistan. Journal of Agricultural Economics, 66(1):66– 86.

Adrakey, K., Malembic-Maher, S., Rusch, A., Ay, J., Riley, L., Ramalanjaona, L., and Fabre, F. (2022). Field and landscape risk factors impacting flavescence doŕee infection : insights from spatial bayesian modelling in the bordeaux vineyards. Phytopathology.

Agrawal, A. and Lively, C. M. (2002). Infection genetics: gene-for-gene versus matching-alleles models and all points in between. Evolutionary Ecology Research, 4(1):91–107.

Ambec, S. and Desquilbet, M. (2012). Regulation of a spatial externality: Refuges versus tax for managing pest resistance. Environmental and resource economics, 51(1):79–104.

Bahri, B., Kaltz, O., Leconte, M., de Vallavieille-Pope, C., and Enjalbert, J. (2009). Tracking costs of virulence in natural populations of the wheat pathogen, Puccinia striiformis f.sp.tritici. BMC Evolutionary Biology, 9(1):26.

Borg, J., Kiær, L. P., Lecarpentier, C., Goldringer, I., Gauffreteau, A., Saint-Jean, S., Barot, S., and Enjalbert, J. (2018). Unfolding the potential of wheat cultivar mixtures: A meta-analysis perspective and identification of knowledge gaps. Field Crops Research, 221:298–313.

Boso, S. and Kassemeyer, H. H. (2008). Different susceptibility of European grapevine cultivars for downy mildew. Vitis, 47(1):39–49.

Bourguet, D., Desquilbet, M., and Lemaríe, S. (2005). Regulating insect resistance management: the case of non-bt corn refuges in the us. Journal of environmental management, 76(3):210–220.

Bove, F., Bavaresco, L., Caffi, T., and Rossi, V. (2019). Assessment of Resistance Components for Improved Phenotyping of Grapevine Varieties Resistant to Downy Mildew. Frontiers in Plant Science, 10(November):1–10.

Bove, F. and Rossi, V. (2020). Components of partial resistance to Plasmopara viticola enable complete phenotypic characterization of grapevine varieties. Scientific Reports, pages 1–12.

Brown, J. K. (2015). Durable Resistance of Crops to Disease: A Darwinian Perspective. Annual Review of Phytopathology, 53:513–539.

Brown, Z. S. (2018). Voluntary Programs To Encourage Refuges for Pesticide Resistance Management: Lessons from a Quasi-Experiment. American Journal of Agricultural Economics, 100(3):844–867. eprint: https://onlinelibrary.wiley.com/doi/pdf/10.1093/ajae/aay004.

Burdon, J. J., Barrett, L. G., Rebetzke, G., and Thrall, P. H. (2014). Guiding deployment of resistance in cereals using evolutionary principles. Evolutionary Applications, 7(6):609–624.

Clin, P., Grognard, F., Andrivon, D., Mailleret, L., and Hamelin, F. M. (2022). Host mixtures for plant disease control: Benefits from pathogen selection and immune priming. Evolutionary Applications, 15(6):967–975.

Clin, P., Grognard, F., Mailleret, L., Val, F., Andrivon, D., and Hamelin, F. M. (2021). Taking advantage of pathogen diversity and immune priming to minimize disease prevalence in host mixtures: a model. Phytopathology®, 111(7):1219–1227.

Day, T. and Gandon, S. (2012). The evolutionary epidemiology of multilocus drug resistance. Evolution; International Journal of Organic Evolution, 66(5):1582–1597.

Djidjou-Demasse, R., Ducrot, A., and Fabre, F. (2017a). Steady state concentration for a phenotypic structured problem modeling the evolutionary epidemiology of spore producing pathogens. Mathematical Models and Methods in Applied Sciences, pages 1–42.

Djidjou-Demasse, R., Moury, B., and Fabre, F. (2017b). Mosaics often outperform pyramids: Insights from a model comparing strategies for the deployment of plant resistance genes against viruses in agricultural landscapes. New Phytologist, pages 239–253.

Fabre, F., Bruchou, C., Palloix, A., and Moury, B. (2009). Key determinants of resistance durability to plant viruses: Insights from a model linking within- and between-host dynamics. Virus research, 141(2):140–149.

Fabre, F., Burie, J.-B., Ducrot, A., Lion, S., Richard, Q., and Djidjou-Demasse, R. (2022). An epi-evolutionary model for predicting the adaptation of spore-producing pathogens to quantitative resistance in heterogeneous environments. Evolutionary Applications, 15(1):95–110. eprint: https://onlinelibrary.wiley.com/doi/pdf/10.1111/eva.13328.

Fabre, F., Rousseau, E., Mailleret, L., and Moury, B. (2015). Epidemiological and evolutionary management of plant resistance: optimizing the deployment of cultivar mixtures in time and space in agricultural landscapes. Evolutionary Applications, 8(10):919–932.

Fischer, A. P., Klooster, A., and Cirhigiri, L. (2019). Cross-boundary cooperation for landscape management: Collective action and social exchange among individual private forest landowners. Landscape and Urban Planning, 188:151–162.

Fouillet, E., Deliére, L., Chartier, N., Munier-Jolain, N., Cortel, S., Rapidel, B., and Merot, A. (2022). Reducing pesticide use in vineyards. Evidence from the analysis of the French DEPHY network. European Journal of Agronomy, 136:126503.

Frantzen, J. and Van den Bosch, F. (2000). Spread of organisms: can travelling and dispersive waves be distinguished? Basic and Applied Ecology, 1(1):83– 92.

Garćıa-Arenal, F. and McDonald, B. A. (2003). An analysis of the durability of resistance to plant viruses. Phytopathology, 93(8):941–952.

Garry, G., Forbes, G., Salas, A., Santa Cruz, M., Perez, W., and Nelson, R. (2005). Genetic diversity and host differentiation among isolates of phytophthora infestans from cultivated potato and wild solanaceous hosts in peru. Plant Pathology, 54(6):740–748.

Geffersa, A., Burdon, J., Macfadyen, S., Thrall, P., Sprague, S., and Barrett, L. (2023). The socio-economic challenges of managing pathogen evolution in agriculture. Philosophical Transactions of the Royal Society B, 378(1873):20220012.

Gessler, C., Pertot, I., and Perazzolli, M. (2011). Plasmopara viticola: A review of knowledge on downy mildew of grapevine and effective disease management. Phytopathologia Mediterranea, 50(1):3–44.

Grosdidier, M., Ioos, R., Husson, C., Cael, O., Scordia, T., and Marcais, B. (2018). Tracking the invasion: dispersal of hymenoscyphus fraxineus airborne inoculum at different scales. FEMS microbiology ecology, 94(5):fiy049.

Kahindo, S. and Blancard, S. (2022). Reducing pesticide use through optimal reallocation at different spatial scales: The case of french arable farming. Agricultural Economics, 53(4):648–666.

Lavigne, F., Martin, G., Anciaux, Y., Papaix, J., and Roques, L. (2020). When sinks become sources: adaptive colonization in asexuals. Evolution, 74(1):29– 42.

Lazarus, W. F. and Dixon, B. L. (1984). Agricultural pests as common property: Control of the corn rootworm. American Journal of Agricultural Economics, 66(4):456–465.

Leach, J. E., Vera Cruz, C. M., Bai, J., and Leung, H. (2001). Pathogen fitness penalty as a predictor of durability of disease resistance genes. Annual review of phytopathology, 39(1):187–224.

Lebeda, A., Petrželová, I., and Maryška, Z. (2008). Structure and variation in the wild-plant pathosystem: Lactuca serriola–bremia lactucae. European Journal of Plant Pathology, 122:127–146.

Lemaríe, S. and Marcoul, P. (2018). Coordination and information sharing about pest resistance. Journal of Environmental Economics and Management, 87:135–149.

Leonard, K. (1977). Selection pressures and plant pathogens. Annals of the New York Academy of Sciences, 287(1):207–222.

Leroy, T., Le Cam, B., and Lemaire, C. (2014). When virulence originates from non-agricultural hosts: New insights into plant breeding. Infection, Genetics and Evolution, 27:521–529.

Leventon, J., Schaal, T., Velten, S., Dänhardt, J., Fischer, J., Abson, D. J., and Newig, J. (2017). Collaboration or fragmentation? biodiversity management through the common agricultural policy. Land use policy, 64:1–12.

Lo Iacono, G., van den Bosch, F., and Gilligan, C. A. (2013). Durable Resistance to Crop Pathogens: An Epidemiological Framework to Predict Risk under Uncertainty. PLoS Computational Biology, 9(1):11–19.

Lo Iacono, G., van den Bosch, F., and Paveley, N. (2012). The evolution of plant pathogens in response to host resistance: Factors affecting the gain from deployment of qualitative and quantitative resistance. Journal of Theoretical Biology, 304:152–163.

Lof, M. E., De Vallavieille-Pope, C., and Van Der Werf, W. (2017). Achieving durable resistance against plant diseases: Scenario analyses with a national-scale spatially explicit model for a wind-dispersed plant pathogen. Phytopathology, 107(5):580–589.

Lof, M. E. and van der Werf, W. (2017). Modelling the effect of gene deployment strategies on durability of plant resistance under selection. Crop Protection, 97:10–17.

McDonald, B. A. and Linde, C. (2002). Pathogen population genetics, evolutionary potential, and durable resistance. Annual Review of Phytopathology, 40:349–379.

Merdinoglu, D., Schneider, C., Prado, E., Wiedemann-Merdinoglu, S., and Mestre, P. (2018). Breeding for durable resistance to downy and powdery mildew in grapevine. OENO One, 52(3):203–209. Number: 3.

Mikaberidze, A., Mundt, C. C., and Bonhoeffer, S. (2015). Invasiveness of plant pathogens depends on the spatial scale of host distribution. Ecological Applications, 26(4):1238–1248.

Mundt, C. C. (1990). Probability of Mutation to Multiple Virulence and Durability of Resistance Gene Pyramids. Phytopathology, 80(2):221–223.

Mundt, C. C. (1991). Probability of Mutation to Multiple Virulence and Durability of Resistance Gene Pyramids: Further Comments. Phytopathology, 81(3):240–242.

Mundt, C. C. (2018). Pyramiding for resistance durability: theory and practice. Phytopathology, 108(7):792–802.

Niks, R. E., Qi, X., Marcel, T. C., et al. (2015). Quantitative resistance to biotrophic filamentous plant pathogens: concepts, misconceptions, and mechanisms. Annu. Rev. Phytopathol, 53(1):10–1146.

Nilusmas, S., Mercat, M., Perrot, T., Djian-Caporalino, C., Castagnone-Sereno, P., Touzeau, S., Calcagno, V., and Mailleret, L. (2020). Multi-seasonal modelling of plant-nematode interactions reveals efficient plant resistance deployment strategies. Evolutionary applications, 13(9):2206–2221.

Ohtsuki, A. and Sasaki, A. (2006). Epidemiology and disease-control under gene-for-gene plant-pathogen interaction. Journal of Theoretical Biology, 238(4):780–794.

Pacilly, F. C., Hofstede, G. J., Lammerts van Bueren, E. T., and Groot, J. C. (2019). Analysing social-ecological interactions in disease control: An agent-based model on farmers’ decision making and potato late blight dynamics. Environmental Modelling and Software, 119(July):354–373.

Pacilly, F. C., Hofstede, G. J., Lammerts van Bueren, E. T., Kessel, G. J., and Groot, J. C. (2018). Simulating crop-disease interactions in agricultural landscapes to analyse the effectiveness of host resistance in disease control: The case of potato late blight. Ecological Modelling, 378(March):1–12.

Papaïx, J., Touzeau, S., Monod, H., and Lannou, C. (2014). Can epidemic control be achieved by altering landscape connectivity in agricultural systems? Ecological Modelling, 284:35–47.

Parlevliet, J. E. (2002). Durability of resistance against fungal, bacterial and viral pathogens; present situation. Euphytica, 124(2):147–156.

Pimentel, D., Acquay, H., Biltonen, M., Rice, P., Silva, M., Nelson, J., Lipner, V., Giordano, S., Horowitz, A., and D’amore, M. (1992). Environmental and economic costs of pesticide use. BioScience, 42(10):750–760.

Pink, D. A. (2002). Strategies using genes for non-durable disease resistance. Euphytica, 124:227–236.

Pirrello, C., Magon, G., Palumbo, F., Farinati, S., Lucchin, M., Barcaccia, G., and Vannozzi, A. (2023). Past, present, and future of genetic strategies to control tolerance to the main fungal and oomycete pathogens of grapevine. Journal of Experimental Botany, 74(5):1309–1330.

R Core Team (2021). R: A Language and Environment for Statistical Computing. R Foundation for Statistical Computing, Vienna, Austria.

Rimbaud, L., Fabre, F., Papaïx, J., Moury, B., Lannou, C., Barret, L. G., and Thrall, P. H. (2021). Models of plant resistance deployment. Annual Review of Phytopathology.

Rimbaud, L., Papaïx, J., Barrett, L. G., Burdon, J. J., and Thrall, P. H. (2018a). Mosaics, mixtures, rotations or pyramiding: What is the optimal strategy to deploy major gene resistance? Evolutionary Applications, 11(10):1791–1810.

Rimbaud, L., Papaïx, J., Rey, J.-F., Barrett, L. G., and Thrall, P. H. (2018b). Assessing the durability and efficiency of landscape-based strategies to deploy plant resistance to pathogens. PLoS computational biology, 14(4):e1006067.

Rimbaud, L., Papaïx, J., Rey, J.-F., Zaffaroni, M., and Gaussen, J.-L. (2023). landsepi: Landscape Epidemiology and Evolution. R package version 1.3.0.

Rousseau, E., Bonneault, M., Fabre, F., Moury, B., Mailleret, L., and Grognard, F. (2019). Virus epidemics, plant-controlled population bottlenecks and the durability of plant resistance. Philosophical Transactions of the Royal Society B: Biological Sciences, 374(1775).

Sacristán, S. and García-Arenal, F. (2008). The evolution of virulence and pathogenicity in plant pathogen populations. Molecular Plant Pathology, 9(3):369–384.

Sapoukhina, N., Durel, C. E., and Le Cam, B. (2009). Spatial deployment of gene-for-gene resistance governs evolution and spread of pathogen populations. Theoretical Ecology, 2:229–238.

Sirami, C., Gross, N., Baillod, A. B., Bertrand, C., Carrié, R., Hass, A., Henckel, L., Miguet, P., Vuillot, C., Alignier, A., Girard, J., Batáry, P., Clough, Y., Violle, C., Giralt, D., Bota, G., Badenhausser, I., Lefebvre, G., Gauffre, B., Vialatte, A., Calatayud, F., Gil-Tena, A., Tischendorf, L., Mitchell, S., Lind-say, K., Georges, R., Hilaire, S., Recasens, J., Solé-Senan, X. O., Roblenõ, I., Bosch, J., Barrientos, J. A., Ricarte, A., Marcos-Garcia, M., Minãno, J., Mathevet, R., Gibon, A., Baudry, J., Balent, G., Poulin, B., Burel, F., Tscharntke, T., Bretagnolle, V., Siriwardena, G., Ouin, A., Brotons, L., Martin, J.-L., and Fahrig, L. (2019). Increasing crop heterogeneity enhances multitrophic diversity across agricultural regions. Proceedings of the National Academy of Sciences, 116(33):16442–16447.

Smulders, M. J., van de Wiel, C. C., and Lotz, L. A. (2021). The use of intellectual property systems in plant breeding for ensuring deployment of good agricultural practices. Agronomy, 11(6):1163.

Tscharntke, T., Grass, I., Wanger, T. C., Westphal, C., and Batáry, P. (2021). Beyond organic farming – harnessing biodiversity-friendly landscapes. Trends in Ecology & Evolution, 36(10):919–930. Publisher: Elsevier.

Vacher, C., Bourguet, D., Desquilbet, M., Lemarié, S., Ambec, S., and Hochberg, M. E. (2006). Fees or refuges: which is better for the sustainable management of insect resistance to transgenic Bt corn? Biology Letters, 2:198–202. 2.

Van Den Bosch, F. and Gilligan, C. A. (2003). Measures of durability of resistance. Phytopathology, 93(5):616–625.

Velten, S., Jager, N. W., and Newig, J. (2021). Success of collaboration for sustainable agriculture: a case study meta-analysis. *Environment*, Development and Sustainability, pages 1–23.

Watkinson-Powell, B., Gilligan, C. A., and Cunniffe, N. J. (2020). When does spatial diversification usefully maximize the durability of crop disease resistance? Phytopathology, 110(11):1808–1820.

Wickham, H., Chang, W., Henry, L., Pedersen, T. L., Takahashi, K., Wilke, C., Woo, K., Yutani, H., Dunnington, D., and RStudio (2022). Create Elegant Data Visualisations Using the Grammar of Graphics. R package version 3.3.6.

Wingen, L., Shaw, M., and Brown, J. (2013). Long-distance dispersal and its influence on adaptation to host resistance in a heterogeneous landscape. Plant Pathology, 62(1):9–20.

Wong, F. P., Burr, H. N., and Wilcox, W. F. (2001). Heterothallism in Plasmopara viticola. Plant Pathology, 50:427–432.

Zaffaroni, M., Rimbaud, L., Rey, J.-F., Papaïx, J., and Fabre, F. (2023). Effects of pathogen reproduction system on the evolutionary and epidemiological control provided by deployment strategies for two major resistance genes in agricultural landscapes. Evolutionary Applications.

Zhan, J., Thrall, P. H., Papaïx, J., Xie, L., and Burdon, J. J. (2015). Playing on a Pathogen’s Weakness: Using Evolution to Guide Sustainable Plant Disease Control Strategies. Annual Review of Phytopathology, 53:19–43.

