## Supporting Information for "Combining single-gene-resistant and pyramided cultivars in agricultural landscape compromises the benefits of pyramiding in most, but not all, productions situations"

### Note S1 Figures

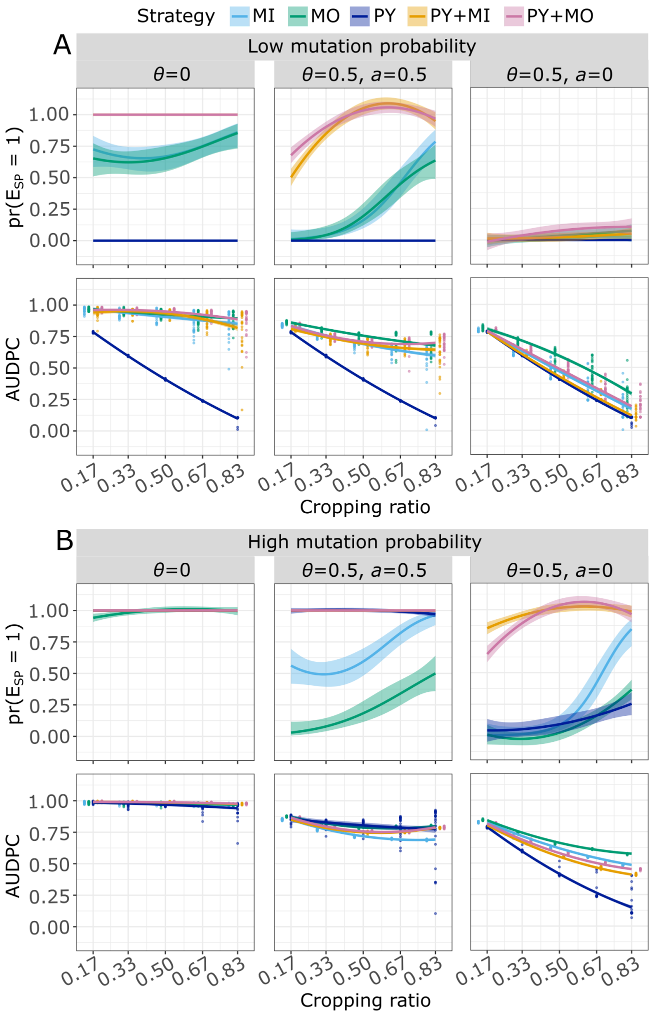


Figure S1: Probability of SP establishment (first row of each panel) and AUDPC (second row of each panel) at low (A, $\tau = 10^{-7}$) and high (B, $\tau = 10^{-4}$) mutation probability, with no fitness cost ($\theta = 0$), with fitness cost for unnecessary virulence ($\theta = 0.5$ and $a = 0.5$), or with fitness cost experienced on all hosts ($\theta = 0.5$ and $a = 0$). Simulations were run with the landscape with 2500 small fields. Panels show the effect on the probability of $E_{SP}$ and AUDPC as a function of the cropping ratio for the five resistant deployment strategies considered. Curves are based on the fitting of second-order logistic (first row) or polynomial (second row) regressions to simulation outputs (represented by points, note that, in the first row, we omitted them, being either 0 or 1); shaded envelopes delimited the 5th and 95th percentiles. When the curves or the shaded envelopes extend beyond the 0-1 range for the probability of $E_{SP}$, it indicates that fitting a second-order logistic regression was impossible, thus a second-order polynomial regression was fitted instead. When the curve for PY+MO is not visible, it overlaps with that of PY+MI.

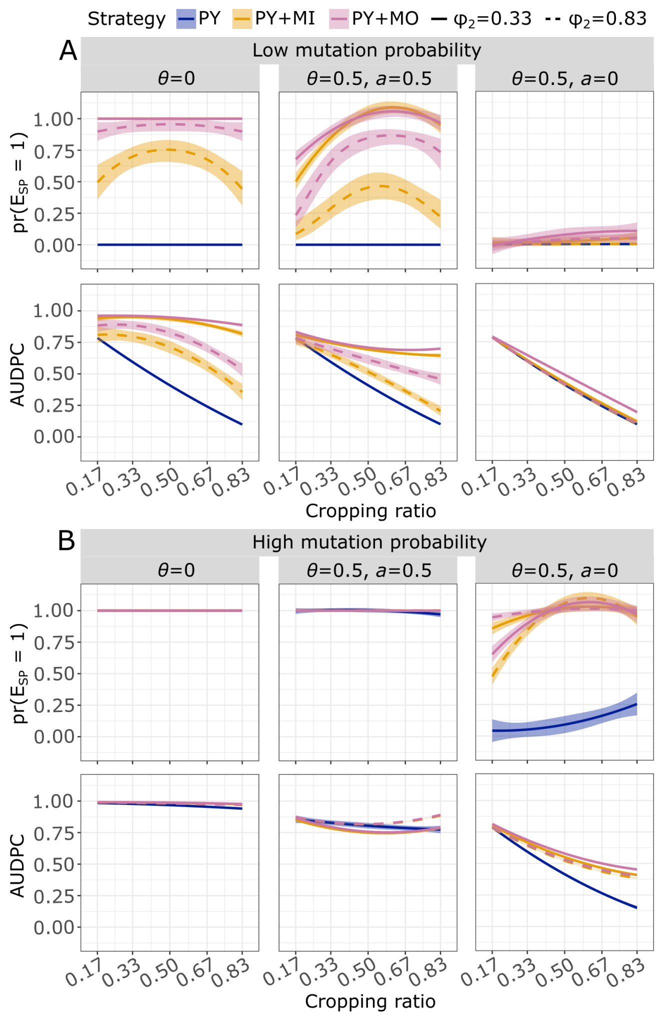


Figure S2: Probability of SP establishment (first row of each panel) and AUDPC (second row of each panel) at low (A, $\tau = 10^{-7}$) and high (B, $\tau = 10^{-4}$) mutation probability, with no fitness cost ($\theta = 0$), with fitness cost for unnecessary virulence ($\theta = 0.5$ and $a = 0.5$), or with fitness cost experienced on all hosts ($\theta = 0.5$ and $a = 0$). Simulations were run with the landscape with 2500 small fields. Panels show the effect on the probability of $E_{SP}$ and AUDPC as a function of the cropping ratio for pyramiding, and for the hybrid strategies, at different proportions of the pyramided cultivars respect to single-gene-resistant cultivars ($\varphi_2$). Curves are based on the fitting of second-order logistic (first row) or polynomial (second row) regressions to simulation outputs; shaded envelopes delimited by the 5th and 95th percentiles. When the curves or the shaded envelopes extend beyond the 0-1 range for the probability of $E_{SP}$, it indicates that fitting a second-order logistic regression was impossible, thus a second-order polynomial regression was fitted instead.

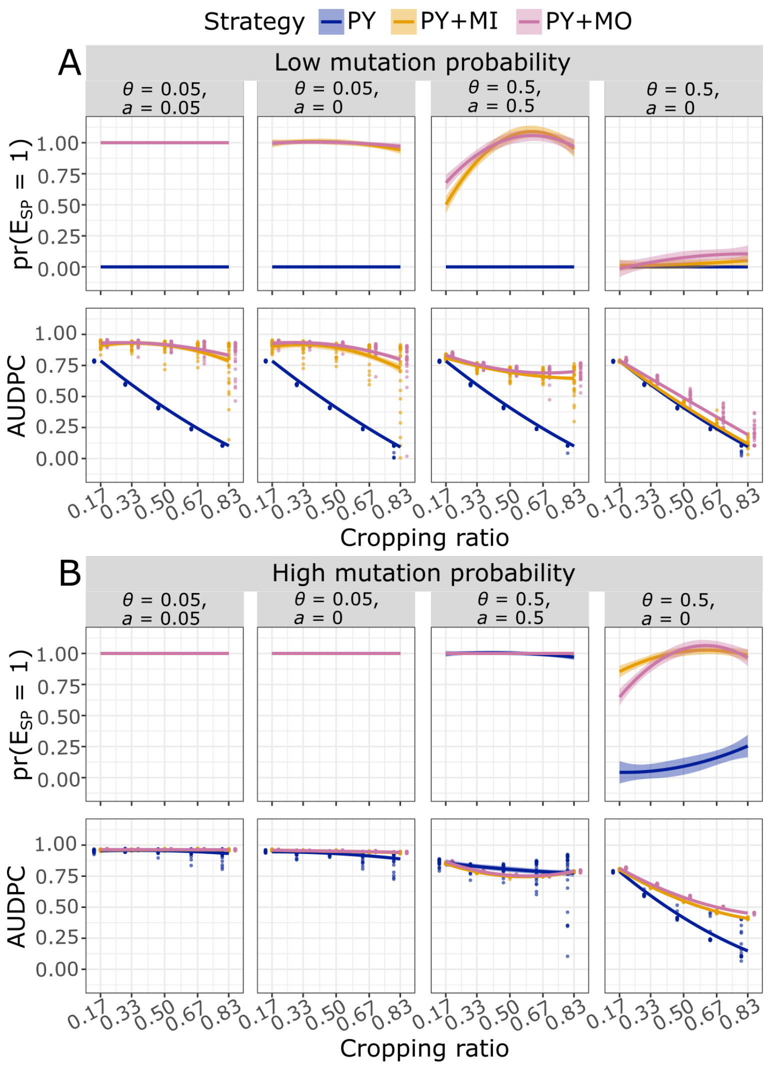


Figure S3: Probability of SP establishment (first row of each panel) and AUDPC (second row of each panel) at low (A, $\tau = 10^{-7}$) and high (B, $\tau = 10^{-4}$) mutation probability, with intermediate or high fitness cost (resp. $\theta = 0.05$ and $\theta = 0.5$), which is paid for unnecessary virulence (resp. $a = 0.05$ and $a = 0.5$), or on any host ($a = 0$). Simulations were run with the landscape with 2500 small fields. Panels show the effect on the probability of $E_{SP}$ and AUDPC as a function of the cropping ratio for pyramiding and for the hybrid strategies. Curves are based on the fitting of second-order logistic (first row) or polynomial regressions (second row) to simulation outputs; shaded envelopes delimited by the 5th and 95th percentiles. When the curves or the shaded envelopes extend beyond the 0-1 range for the probability of $E_{SP}$, it indicates that fitting a second-order logistic regression was impossible, thus a second-order polynomial regression was fitted instead. When the curve for PY+MO is not visible, it overlaps with that of PY+MI.

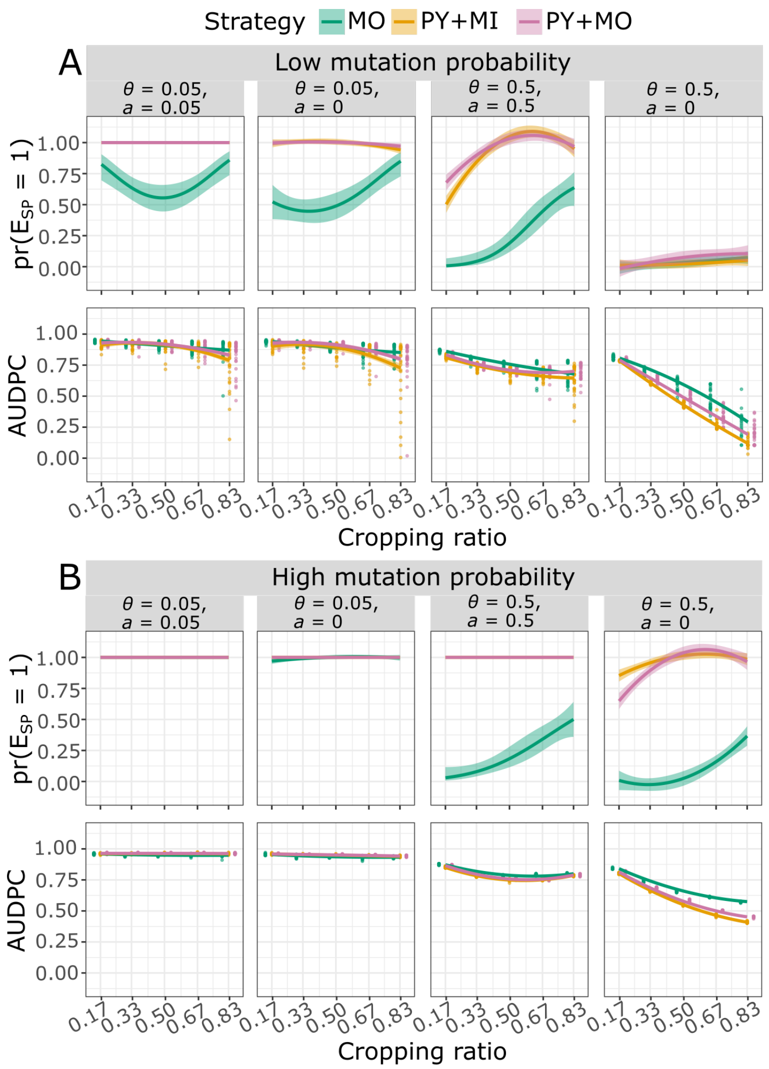


Figure S4: Probability of SP establishment (first row of each panel) and AUDPC (second row of each panel) at low (A, $\tau = 10^{-7}$) and high (B, $\tau = 10^{-4}$) mutation probability, with intermediate or high fitness cost (resp. $\theta = 0.05$ and $\theta = 0.5$), which is paid for unnecessary virulence (resp. $a = 0.05$ and $a = 0.5$), or on any host ($a = 0$). Simulations were run with the landscape with 2500 small fields. Panels show the effect on the probability of $E_{SP}$ and AUDPC as a function of the cropping ratio for mosaic and for the hybrid strategies. Curves are based on the fitting of second-order logistic (first row) or polynomial regressions (second row) to simulation outputs; shaded envelopes delimited by the 5th and 95th percentiles. When the curves or the shaded envelopes extend beyond the 0-1 range for the probability of $E_{SP}$, it indicates that fitting a second-order logistic regression was impossible, thus a second-order polynomial regression was fitted instead. When the curve for PY+MO is not visible, it overlaps with that of PY+MI.

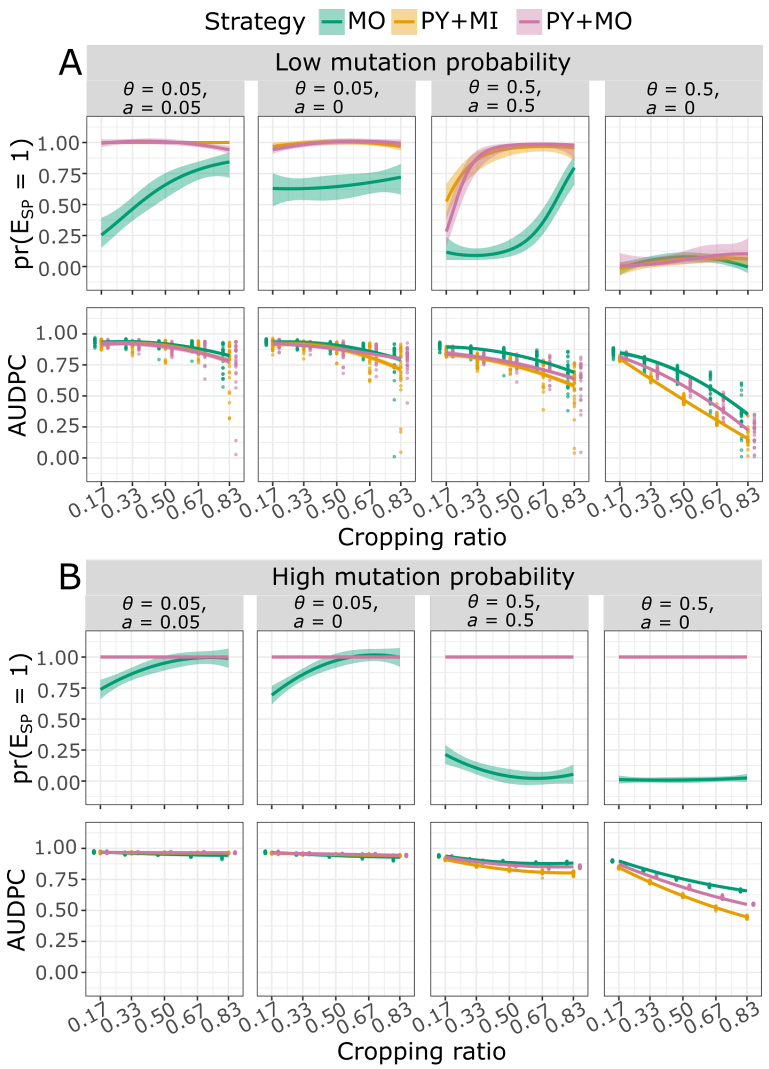


Figure S5: Probability of SP establishment (first row of each panel) and AUDPC (second row of each panel) at low (A, $\tau = 10^{-7}$) and high (B, $\tau = 10^{-4}$) mutation probability, with intermediate or high fitness cost (resp. $\theta = 0.05$ and $\theta = 0.5$), which is paid for unnecessary virulence (resp. $a = 0.05$ and $a = 0.5$), or on any host ($a = 0$). Simulations were run with the landscape with 100 large fields. Panels show the effect on the probability of $E_{SP}$ and AUDPC as a function of the cropping ratio for mosaic and for the hybrid strategies. Curves are based on the fitting of second-order logistic (first row) or polynomial regressions (second row) to simulation outputs; shaded envelopes delimited by the 5th and 95th percentiles. When the curves or the shaded envelopes extend beyond the 0-1 range for the probability of $E_{SP}$, it indicates that fitting a second-order logistic regression was impossible, thus a second-order polynomial regression was fitted instead. When the curve for PY+MO is not visible, it overlaps with that of PY+MI.

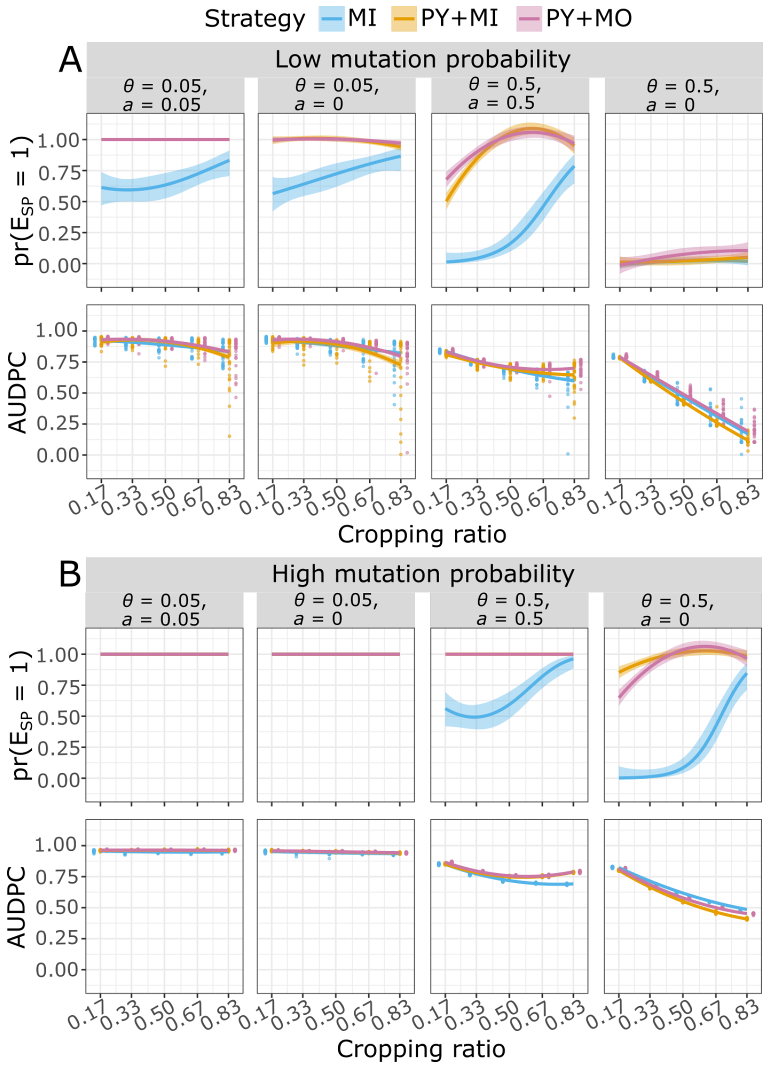


Figure S6: Probability of SP establishment (first row of each panel) and AUDPC (second row of each panel) at low (A, $\tau = 10^{-7}$) and high (B, $\tau = 10^{-4}$) mutation probability, with intermediate or high fitness cost (resp. $\theta = 0.05$ and $\theta = 0.5$), which is paid for unnecessary virulence (resp. $a = 0.05$ and $a = 0.5$), or on any host ($a = 0$). Simulations were run with the landscape with 2500 small fields. Panels show the effect on the probability of $E_{SP}$ and AUDPC as a function of the cropping ratio for mixture and for the hybrid strategies. Curves are based on the fitting of second-order logistic (first row) or polynomial regressions (second row) to simulation outputs; shaded envelopes delimited by the 5th and 95th percentiles. When the curves or the shaded envelopes extend beyond the 0-1 range for the probability of $E_{SP}$, it indicates that fitting a second-order logistic regression was impossible, thus a second-order polynomial regression was fitted instead. When the curve for PY+MO is not visible, it overlaps with that of PY+MI.

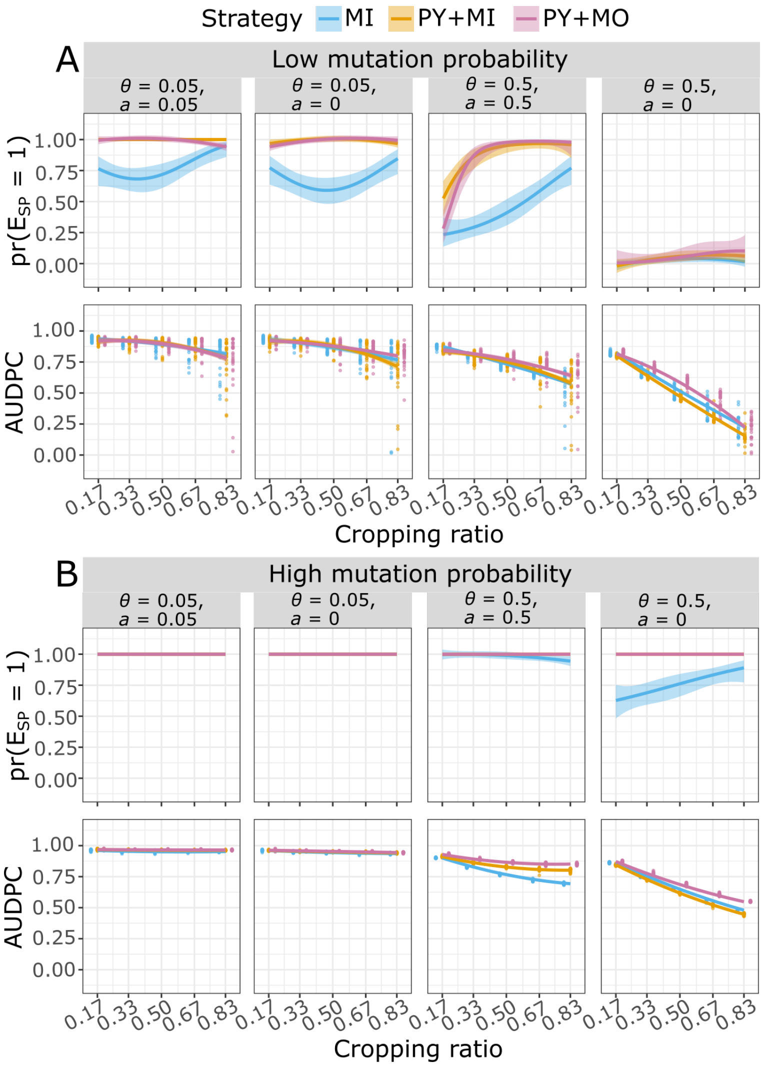


Figure S7: Probability of SP establishment (first row of each panel) and AUDPC (second row of each panel) at low (A, $\tau = 10^{-7}$) and high (B, $\tau = 10^{-4}$) mutation probability, with intermediate or high fitness cost (resp. $\theta = 0.05$ and $\theta = 0.5$), which is paid for unnecessary virulence (resp. $a = 0.05$ and $a = 0.5$), or on any host ($a = 0$). Simulations were run with the landscape with 100 large fields. Panels show the effect on the probability of $E_{SP}$ and AUDPC as a function of the cropping ratio for mixture and for the hybrid strategies. Curves are based on the fitting of second-order logistic (first row) or polynomial regressions (second row) to simulation outputs; shaded envelopes delimited by the 5th and 95th percentiles. When the curves or the shaded envelopes extend beyond the 0-1 range for the probability of $E_{SP}$, it indicates that fitting a second-order logistic regression was impossible, thus a second-order polynomial regression was fitted instead. When the curve for PY+MO is not visible, it overlaps with that of PY+MI.

### Note S2 Model equations

In the present work, simulations are run with the model presented in Rimbaud et al. (2018) and updated in?. Simulations are splitted into two distinct time periods: *i*) within cropping season, where multiple clonal reproduction events take place, and *ii*) between cropping seasons when a single sexual reproduction event takes place. The entire model is described in the following sections. Note that in the equations below only major resistance genes are considered. See Rimbaud et al. (2018) for details on model equations considering both major genes and quantitative resistance traits.

#### S2. 1 Host and pathogen demo-genetic dynamics within cropping seasons

The demo-genetic dynamics of the host-pathogen interaction are based on a HLIR structure ("healthy-latent-infectious-removed"). Thus, in the following, $H_{i,v,t}$, $L_{i,v,p,t}$, $I_{i,v,p,t}$, $R_{i,v,p,t}$, and $Pr_{i,p,t}$ respectively denote the number of healthy, latent, infectious and removed individuals (in this model, an "individual" is a given amount of plant tissue, and is referred to as a "host" hereafter for simplicity), and pathogen propagules in field $i$ ($i$=1,...,$J$), for cultivar $v$ ($v$=1,...,$V$), pathogen genotype $p$ ($p$=1,...,$P$) at time step $t$ ($t$=1,...,$T \times Y$). $T$ is the number of time steps in a cropping season and $Y$ the number of simulated years (*i.e.* cropping seasons). Since the host is cultivated, we assume there is no host reproduction, dispersal or natural mortality (leaf senescence near the end of the cropping season is considered as part of host harvest).

**Host growth.** Only healthy hosts (denoted as $H_{i,v,t}$) are assumed to contribute to growth of the crop. Thus, at each step $t$ during a cropping season, the plant cover of cultivar $v$ in field $i$ increases as a logistic function, and the new amount of healthy plant tissue is:

$$H_{i,v,t+1} = H_{i,v,t}\left[1 + \delta_v \times \left(1 - \frac{N_{i,v,t}}{K_{i,v}}\right)\right] \tag{1}$$

with $\delta_v$ the growth rate of cultivar $v$; $N_{i,v,t} = H_{i,v,t} + \sum_{p=1}^{P}\left(L_{i,v,p,t} + I_{i,v,p,t} + R_{i,v,p,t}\right)$ the total number of hosts in field $i$ for cultivar $v$ and at time $t$; and $K_{i,v} = A_i \times C_v^{max}$ the carrying capacity of cultivar $v$ in field $i$, which depends on $A_i$, the area of the field, and $C_v^{max}$, the maximal density for cultivar $v$. Note that equation (1) adequately approximates a continuous time logistic function only when $\delta_v \leq 1$, otherwise negative recruitment could be generated. When a mixture of several cultivars is present in the same field, decreased growth due to susceptible plants being diseased is not compensated by increased growth of resistant plants.

**Contamination of healthy hosts.** The healthy compartment ($H$) is composed of hosts which are free of pathogen propagules ($H^1$), as well as hosts contaminated (but not yet infected) by the arrival of such propagules ($H^2$). At the beginning of each step, all healthy hosts are considered free of propagules ($H^1$). Then at time $t$ in field $i$ and for cultivar $v$, the number of contaminable hosts (*i.e.* accessible to pathogen propagules, denoted as $H_{i,v,t}^{contaminable}$) depends on the proportion of healthy hosts ($H_{i,v,t}^1$) in the host population ($N_{i,v,t}$):

$$H_{i,v,t}^{contaminable} \sim Binomial\left(H_{i,v,t}^1; \pi\left(\frac{H_{i,v,t}^1}{N_{i,v,t}}\right)\right) \tag{2}$$

with $\pi(x) = \frac{1-e^{-\kappa x^\sigma}}{1-e^{-\kappa}}$, a sigmoid function with $\pi(0) = 0$ and $\pi(1) = 1$, giving the probability for a healthy host to be contaminated. Here, we assume that healthy hosts are not equally likely to be contacted by propagules, for instance because of plant architecture. Moreover, as the local severity of disease increases, eventually the probability for a single propagule to contaminate a healthy host declines due to the decreased availability of host tissue.

Following the arrival of propagules of pathogen genotype $p$ in field $i$ at time $t$ (denoted as $Pr_{i,p,t}^4$, see below), susceptible hosts become contaminated. The pathogen genotypes of these propagules are distributed among contaminable hosts according to their proportional representation in the total pool of propagules. Thus, for cultivar $v$, the vector describing the maximum number of contaminated hosts by each pathogen genotype (denoted as $[H_{i,v,t}^{maxConta}]_{p=1,\ldots,P}$) is given by a multinomial draw:

$$[H_{i,v,t}^{maxConta}]_{p=1,\ldots,P} \sim Multinomial\left(H_{i,v,t}^{contaminable}; \left[\frac{Pr_{i,p,t}^4}{\sum_{p=1}^{P} Pr_{i,p,t}^4}\right]_{p=1,\ldots,P}\right) \tag{3}$$

However, the number of deposited propagules ($Pr_{i,p,t}^4$) may be smaller than the maximal number of contaminated hosts ($H_{i,v,p,t}^{maxConta}$). Thus, the true number of hosts of cultivar $v$, contaminated by pathogen genotype $p$ in field $i$ at $t$ (denoted as $H_{i,v,p,t}^2$) is given by:

$$[H_{i,v,p,t}^1 \rightarrow H_{i,v,p,t}^2] = min\left(H_{i,v,p,t}^{maxConta}; Pr_{i,p,t}^4\right) \tag{4}$$

**Infection.** Between $t$ and $t+1$, in field $i$, contaminated hosts ($H_{i,v,p,t}^2$) become infected (state L) with probability $e_{v,p}$, which depends on the maximum expected infection probability, $e_{max}$, and the interaction between host ($v$) and pathogen ($p$) genotypes:

$$[H_{i,v,p,t}^2 \rightarrow L_{i,v,p,t+1}] \sim Binomial\left(H_{i,v,p,t}^2; e_{v,p}\right) \tag{5}$$

$$e_{v,p} = e_{max} \times \prod_{g=1}^{G} INF_{ig_g(p),mg(v)}^g \tag{6}$$

$INF^g$ represent the infectivity matrix for major gene $g$ which summarizes the possible interactions between potential host resistance genes ($mg$) and associated pathogen infectivity genes ($ig$), see Table 1 in the main text for an example.

**Latent period.** Infected hosts become infectious (state I) after a latent period (LI) drawn from a Gamma distribution (a flexible continuous distribution from which durations in the interval $[0;+\inf[$ can be drawn) parameterised with expected value, $\Gamma_{exp}$, and variance, $\Gamma_{var}$:

$$(LI) \sim Gamma(\Gamma_{exp}; \Gamma_{var}) \tag{7}$$

Note, the usual shape and scale parameters of a Gamma distribution, $\beta_1$ and $\beta_2$, can be calculated from the expectation and variance, exp and var, with: $\beta_1 = \frac{exp^2}{var}$ and $\beta_2 = \frac{var}{exp}$, respectively.

**Infectious period.** Finally, infectious hosts become epidemiologically inactive (*i.e.* they no longer produce propagules, thus are in state R, "removed") after an infectious period (IR) drawn from a Gamma distribution parameterised with expected value, $\Upsilon_{exp}$ and variance, $\Upsilon_{var}$, similar to the latent period:

$$(IR) \sim Gamma(\Upsilon_{exp}; \Upsilon_{var}) \tag{8}$$

**Pathogen clonal reproduction.** In field $i$ at time $t$, infectious hosts associated with pathogen genotype $p$ produce a total number of propagules (denoted as $Pr^1_{i,p,t}$), drawn from a Poisson distribution with parameter $r_{exp}$ corresponding to the expected number of propagules produced by a single infectious host per time step:

$$Pr^1_{i,p,t} \sim Poisson(r_{exp} \sum_{v=1}^{V} I_{i,v,p,t}) \tag{9}$$

**Pathogen mutation.** The following algorithm is repeated independently for every potential infectivity gene g:

1. the pathotype (*i.e.* the level of adaptation with regard to major gene $g$, indexed by $q$; $q = 1, \ldots, Q_g$; with $Q_g = 2$ since the pathotype is either infective, or non-infective) of the pathogen propagules is retrieved from their genotype $p$;

2. propagules can mutate from pathotype $q$ to pathotype $q'$ with probability $m^g_{qq'}$ such as $m^g_{qq'} = \tau_g$ if $q' \neq q$ (hence $m^g_{qq} = 1 - \tau_g$ since $Q_g = 2$). Thus, in field $i$ at time $t$, the vector of the number of propagules of each pathotype arising from pathotype $q$ (denoted as $[M^g_{i,q,q',t}]_{q'=1,\ldots,Q_g}$) is given by a multinomial draw:

$$[M^g_{i,q,q',t}]_{q'=1,\ldots,Q_g} \sim Multinomial\left(Pr^1_{i,p,t}; [m^g_{qq'}]_{q'=1,\ldots,Q_g}\right) \tag{10}$$

3. the total number of propagules belonging to pathotype $q'$ and produced in field $i$ at time $t$ (denoted as $Pr^2_{i,q',t}$) is:

$$Pr^2_{i,q',t} = \sum_{q=1}^{Q_g} M^g_{i,q,q',t} \tag{11}$$

4. the new propagule genotype $p'$ is retrieved from its new pathotype ($q'$), and the number of propagules is incremented using a variable denoted as $Pr^3_{i,p',t}$.

In this model, it should be noted that the mutation probability $\tau_g$ is not the classic mutation rate (*i.e.* the number of genetic mutations per generation per base pair), but the probability for a propagule to change its infectivity on a resistant cultivar carrying major gene $g$. This probability depends on the classic mutation rate, the number and nature of the specific genetic mutations required to overcome major gene $g$, and the potential dependency between these mutations.

**Pathogen dispersal.** Propagules (both clonal and sexual, see below) can migrate from field $i$ (whose area is $A_i$) to field $i'$ (whose area is $A'_i$) with probability $\mu_{ii'}$, computed from:

$$\mu_{ii'} = \frac{\int_{A_i}\int_{A_{i'}} g(\|z'-z\|)dzdz'}{A_i} \tag{12}$$

with $\|z'-z\|$ the Euclidian distance between locations $z$ and $z'$ in fields $i$ and $i'$, respectively, and $g(\|z'-z\|) = \frac{(\beta-2)(\beta-1)}{2\pi\alpha^2}\left(1+\frac{\|z'-z\|}{\alpha}\right)^{-\beta}$ the two-dimensional power law dispersal kernel of the propagules. The computation of $\mu_{ii'}$ probability is performed using the *CaliFloPP* algorithm (Bouvier et al., 2009). Thus, at time $t$, the vector of the number of propagules of genotype $p$ migrating from field $i$ to each field $i'$ (denoted as $[D_{i,i',p,t}]_{i'=1,\dots,J}$) is:

$$[D_{i,i',p,t}]_{i'=1,\dots,J} \sim Multinomial(Pr^3_{i,p,t}; [\mu_{i,i'}]_{i'=1,\dots,J}) \tag{13}$$

and the total number of propagules arriving in field $i'$ at time $t$ (denoted as $Pr^4_{i',p,t}$) is:

$$Pr^4_{i',p,t} = \sum_{i=1}^{J} D_{i,i',p,t} \tag{14}$$

We consider that propagules landing outside the boundaries of the simulated landscape are lost (absorbing boundary condition), and there are no propagule sources external to the simulated landscape.

#### S2. 2 Host and pathogen demo-genetic dynamics between cropping seasons

**Seasonality.** Let $t^0(y)$ and $t^f(y)$ denote the first and last days of cropping season $y$ ($y = 1, \ldots, Y$), respectively. The plant cover in field $i$ for cultivar $v$ at the beginning of cropping season $y$ is set at $H_{i,v,t^0(y)} = A_i \times C_v^0 \times \mathbb{I}_{v(i)=v}$, with $C_v^0$ the plantation density of cultivar $v$ and $\mathbb{I}_{v(i)}$ an indicative variable set at 1 when field $i$ is cultivated with cultivar $v$ and 0 otherwise. At the end of a cropping season, the host is harvested. We assume that the pathogen needs a green bridge to survive the off-season. This green bridge could, for example, be a wild reservoir or volunteer plants remaining in the field (*e.g.* owing to incomplete harvest or seedlings). The size of this reservoir imposes a bottleneck for the pathogen population. The number of remaining infected hosts in field $i$ for cultivar $v$ and pathogen genotype $p$ (denoted by $I^*_{i,v,p,t^f(y)}$) at the end of the off-season is given by:

$$I^*_{i,v,p,t^f(y)} \sim Binomial(L_{i,v,p,t^f(y)}) + I_{i,v,p,t^f(y)}); \lambda) \quad (15)$$

with $\lambda$ the survival probability of infected hosts. Considering that those hosts produce propagules during their whole infectious period, we compute an equivalent number of infectious hosts by multiplying $I^{eq}_{i,v,p,t^f(y)} = \Upsilon_{exp} \times I^*_{i,v,p,t^f(y)}$. The remaining hosts $I^{eq}_{i,v,p,t^f(y)}$ produce clonal and sexual propagules. Clonal propagules can mutate exactly as happens during the cropping season. The production of propagules through sexual reproduction and the possible genetic recombination are detailed in the following section. Propagules, either clonal or sexual, produced between cropping seasons are uniformly released throughout the following cropping season, constituting the primary inoculum.

**Pathogen sexual reproduction.** In field $i$, the pool of infectious hosts associated to the same cultivar $v$, $I_{i,v,p,t^f(y)}$, undertakes sexual reproduction. Two parental infectious hosts, respectively infected by pathogens $Par_1$ and $Par_2$ are randomly sampled without replacement from the pool of infectious hosts. The couple $c = \{Par_1; Par_2\}$ produce $P^{sex}_{v,c}$ propagules, drawn from a Poisson distribution whose expectation is the sum of the number $r_{exp}$ of propagules produced by each of the parental infectious hosts:

$$P^{sex}_{v,c} \sim Poisson(r_{expv,c} = r_{exp} + r_{exp} = 2 \times r_{exp}) \quad (16)$$

Then the genotype of each propagule is retrieved from parental genotypes: the genotype at every locus $g$ is randomly sampled among the two parents $\{Par_1; Par_2\}$. For example, assuming that parental infection $Par_1$ carries infectivity genes to resistance gene $R_1$ (which corresponds to a genotype "10") and parental infection $Par_2$ carries the infectivity genes to resistance $R_2$ (genotype "01"), the resulting propagule genotype could be either the same as one of the two parents, or a superpathogen genotype "11", or a wild-type genotype "00". This process is iterated for all the pairs $c$ = 1,..., $C$ of infectious hosts associated to all the cultivars $v = 1,..., V$ in a given field $i$, resulting in a total number of sexual propagules:

$$P_i^{sex} = \sum_{v=1}^{V} \sum_{c=1}^{C} P_{v,c}^{sex} \tag{17}$$

#### S2. 3 Initial conditions

At the beginning of a simulation, healthy hosts are planted in each field. The initial pathogen population is assumed to be totally non-adapted to the resistance genes, and is only present in susceptible fields with probability $\Phi$. Then the initial number of infectious hosts in these fields is:

$$I_{i,v=1,p=1,t=1} \sim Binomial(H_{i,v=1,t=1}; \Phi) \tag{18}$$

### References


Bouvier, A., Kiêu, K., Adamczyk, K., and Monod, H. (2009). Computation of the integrated flow of particles between polygons. *Environmental Modelling & Software*, 24(7):843–849.

Rimbaud, L., Papaïx, J., Rey, J.-F., Barrett, L. G., and Thrall, P. H. (2018). Assessing the durability and efficiency of landscape-based strategies to deploy plant resistance to pathogens. *PLoS computational biology*, 14(4):e1006067.
